# Microbial single-cell RNA sequencing by split-pool barcoding

**DOI:** 10.1101/869248

**Authors:** Anna Kuchina, Leandra M. Brettner, Luana Paleologu, Charles M. Roco, Alexander B. Rosenberg, Alberto Carignano, Ryan Kibler, Matthew Hirano, R. William DePaolo, Georg Seelig

## Abstract

Single-cell RNA-sequencing (scRNA-seq) has become an essential tool for characterizing multi-celled eukaryotic systems but current methods are not compatible with bacteria. Here, we introduce microSPLiT, a low cost and high-throughput scRNA-seq method that works for gram-negative and gram-positive bacteria and can resolve transcriptional states that remain hidden at a population level. We applied microSPLiT to >25,000 *Bacillus subtilis* cells sampled from different growth stages, creating a detailed atlas of changes in metabolism and lifestyle. We not only retrieve detailed gene expression profiles associated with known but rare states such as competence and PBSX prophage induction, but also identify novel and unexpected gene expression states including heterogeneous activation of a niche metabolic pathway in a subpopulation of cells. microSPLiT empowers high-throughput analysis of gene expression in complex bacterial communities.

## Main Text

Gene expression in bacteria is highly heterogeneous even in isogenic populations grown under the same lab conditions. Using bet-hedging strategies, bacteria differentiate into subpopulations that assume different roles for the survival of the community (*1*, *2*). For example, gene expression programs governing developmental and stress-response states such as competence or antibiotic resistance may switch on stochastically in a small number of single cells (*3*–*5*). Population level gene expression measurements are insufficient to resolve such rare states which, to date, have been characterized only in tractable model systems and through single-cell methods such as fluorescence microscopy that can only measure a limited set of reporter genes at a time (*6*).

Single-cell RNA-seq (scRNA-seq) methods developed for use with eukaryotic cells can provide comprehensive gene expression profiles for tens of thousands of cells (*7*–*11*). However, although the need for microbial scRNA-seq has been recognized, early attempts have been limited to relatively small cell numbers because of the technical challenges associated with adapting scRNA-seq technology to microbes (Figure 1A)(*12*–*14*). Specifically, bacteria have very low mRNA content, typically about two orders of magnitude less than human cells (*13*). Separation of mRNA from rRNA is challenging because bacterial mRNA is not polyadenylated. Bacteria have a wide diversity of cell walls and membranes which interfere with the lysis or permeabilization steps required for scRNA-seq and their small size can hinder microfluidic single-cell isolation. Here, we present microSPLiT (Microbial Split-Pool Ligation Transcriptomics), a scRNA-seq platform that can overcome these challenges and become a transformative technology for microbiology research.

**Fig. 1.**
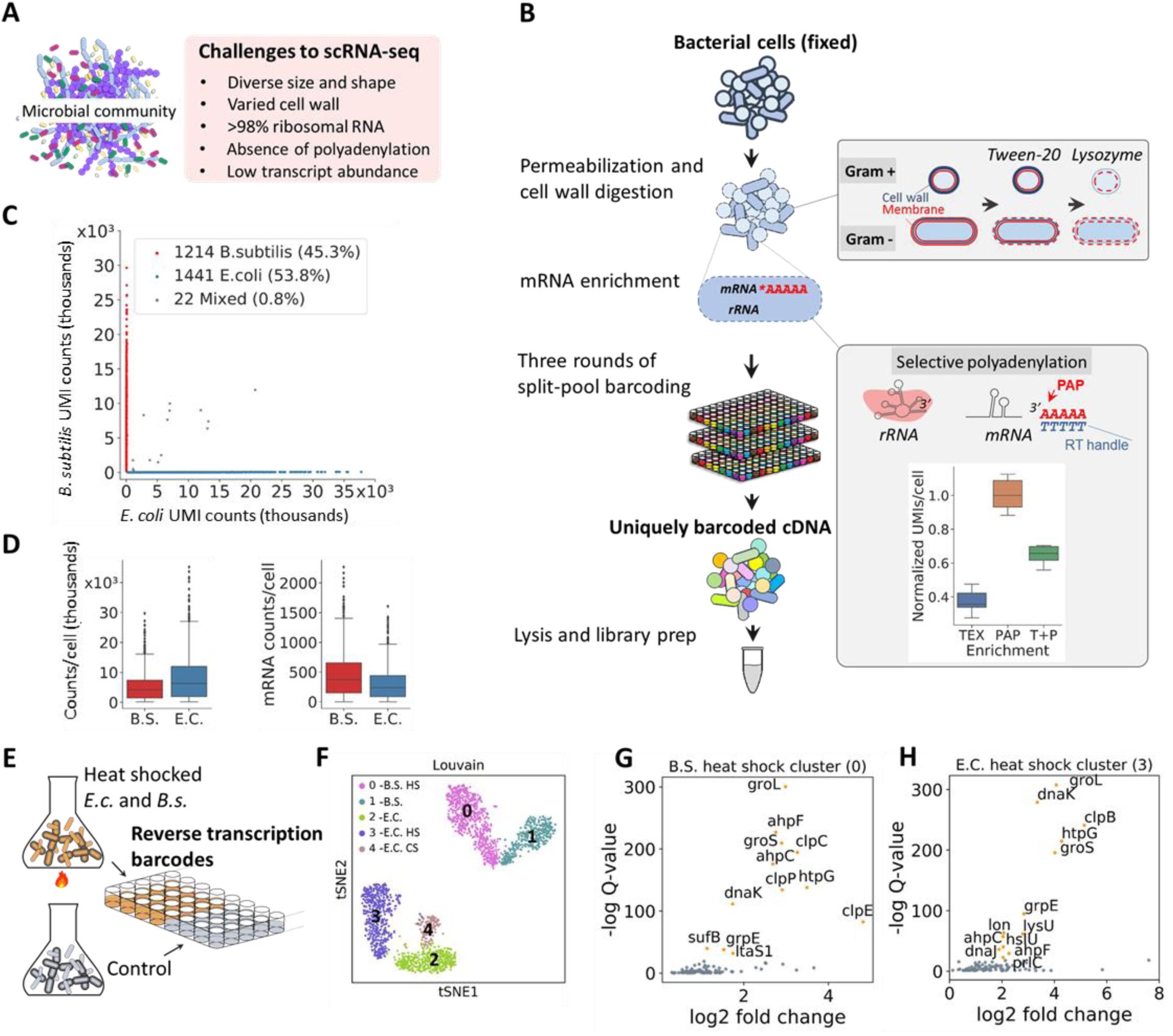
microSPLiT development and validation. **(A)** Challenges associated with single-cell RNA sequencing of bacteria. (**B**) microSPLiT method summary. Fixed gram-positive and gram-negative bacterial cells are permeabilized with a mild detergent Tween-20 and their cell walls are degraded by lysozyme. The mRNA is then polyadenylated in-cell with *E.coli* Poly(A) Polymerase I (PAP). Inset: comparison of mRNA read counts (normalized) obtained from PAP-treated cells compared with cells treated with Terminator 5’ phosphate-dependent exonuclease (TEX) and both methods consecutively (T+P). The cellular RNA then undergoes three rounds of combinatorial barcoding including in-cell reverse transcription (RT) and two in-cell ligation reactions, followed by lysis and library preparation. (**C**) Barnyard plot for the *E. coli* and *B. subtilis* species-mixing experiment. Each dot corresponds to a putative single-cell transcriptome. UMI – unique molecular identifier. (**D**) Total (in thousands) and mRNA UMI counts per cell for both species. (**E**) Summary of the heat shock experiment showing the first barcode as sample identifier. (**F**) t-stochastic neighbor embedding (t-SNE) of the data from heat shock experiment showing distinct clusters. “HS” – heat shocked. (**G**) and (**H**) Genes enriched in the B.S and E.C. heat shock clusters, respectively.

### Developing microSPLiT

SPLiT-seq, which labels the cellular origin of RNA through combinatorial barcoding, provides a starting point for bacterial single cell transcriptomics (*7*). In SPLiT-seq, cells are fixed, permeabilized and mRNA is converted to cDNA through in-cell reverse transcription (RT) with barcoded poly-T and random hexamer primers in a multi-well format. Cells are then pooled, randomly split into a new 96-well plate, and a well-specific barcode is appended to the cDNA through ligation. This split-ligation-pool cycle is repeated and a fourth, optional barcode is added during sequencing library preparation to ensure that each cell acquires a unique barcode combination with very high likelihood (Figure 1B).

Because SPLiT-seq does not require cell isolation, it is compatible with a wide range of cell shapes and sizes. Moreover, because SPLiT-seq already uses random hexamer primers in addition to poly-T primers for RT, we reasoned that it might be suitable for detecting bacterial mRNA. However, a direct application of the mammalian SPLiT-seq protocol to bacteria, not surprisingly, resulted in low total UMI counts (<100 max UMIs/cell, median 0 mRNA reads/cell) and a bias toward gram-negative over gram-positive bacteria (**Figure S1**).

Next, we set out to optimize sample processing steps for bacteria. We took advantage of SPLiT-seq’s multiplexing capabilities to test a wide range of approaches to cell wall removal and membrane permeabilization. We settled on a combination of a mild detergent, Tween-20, and lysozyme, as that treatment protocol demonstrated the best capture for both the gram-positive and gram-negative bacteria tested. (Figure 1B, **inset, and Supplementary Table S1**). Then, we compared different methods that would be compatible for in-cell mRNA enrichment. We tested polyadenylation with *E.coli* Poly(A) Polymerase I (PAP) which was previously used to preferentially polyadenylate mRNA (*15*), 5’-phosphate-dependent exonuclease (“Terminator”, Epicentre) treatment and reverse transcription with ribosomal RNA-specific probes followed by RNaseH-mediated degradation (**Figure S2 and Supplementary Table S1**). We found that the treatment of fixed and permeabilized cells with PAP resulted in the highest (about 2.5-fold, or approximately 7% of total RNA) enrichment of mRNA reads (Figure 1B, **inset, and Figure S2**). We also optimized the fixation protocol as well as the downstream enzymatic reaction conditions (**Supplementary Table S1 and Materials and Methods).** Notably, we found that RT resulted in cell clumping and that mild sonication after this step was necessary to reliably obtain single cell suspensions (**Figure S3**).

### microSPLiT generates high-quality single-cell RNA-seq data

In order to validate microSPLiT performance on a mixture of gram-positive and gram-negative organisms, we grew *Escherichia coli* MW1255 and *Bacillus subtilis* PY79 cells to OD0.5 and subjected half of each culture to a brief 47C heat-shock. We performed a microSPLiT experiment on both samples, using the first barcode as a sample identifier. We prepared and sequenced a cDNA library from 2677 total bacteria from heat-shocked and control treatments and aligned the reads to a combined *B. subtilis*-*E. coli* genome. 99.2% of the putative single cell transcriptomes were unambiguously assigned to a single species (Figure 1C). We sampled a median of 237 unique mRNA transcripts per cell for *E. coli* and 376 for *B. subtilis*, or approximately 5-10% of the estimated total mRNA (*16*). In total, we detected 3717 genes for *E. coli* and 3476 genes for *B. subtilis* (Figure 1D).

Next, we tested whether microSPLiT could detect transcriptional responses to heat shock (Figure 1E). Unsupervised clustering identified five distinct clusters which were visualized by t-distributed stochastic neighbor embedding (t-SNE) (Figure 1F). The first barcode identified two pairs of clusters corresponding to the heat treated and control cultures, and gene expression analysis within each pair further labeled them as corresponding to *B. subtilis* and *E. coli* cells (**Figure S4A**). Analysis of genes enriched within each cluster revealed classical heat shock genes differentially expressed in each of the E.C. and *B. subtilis* heat treated clusters. We detected induction of genes for abundant class I-IV and VI *B. subtilis* heat shock genes such as *groEL*, *dnaK, clpC* and *htpG* operons (Figures 1G, **S4B-C**) as well as both of the major chaperone systems of E.C. (*groEL* and *dnaK* operons, Figures 1H, **S4B-C**) (*17*, *18*).

Unexpectedly, we found an additional small cluster, representing *E. coli* cells from both control and heat treated samples that expressed a dramatically different signature of DEAD-box helicase *deaD* induction as well as cold shock genes *cspA-G* consistent with a transcriptional response to cold (Figures 1F **and S4D**) (*19*). This subpopulation of *E. coli* might be displaying a very rapid response to cold from a brief cold centrifugation step performed as the first step in sample preparation before formaldehyde fixation.

### Transcriptional patterns during *B. subtilis* growth in rich medium

Next, we applied microSPLiT to capture transcriptional states across the *B. subtilis* growth curve in a rich medium (LB). In total, we sampled ten optical density (OD) points along the growth curve of the laboratory strain PY79 ranging from OD 0.5 (early exponential phase) to 6.0 (early stationary phase). Four of the OD points were sampled in both replicates of microSPLiT while the rest were sampled only in one of the experiments (Figure 2A). In both experiments, the first barcode was used to record sample identity (i.e. OD). The data from two different experiments are consistent and produced a combined dataset of 25,214 cells (Figures 2B, **S5 and S6**). Unsupervised clustering of the combined datasets revealed 14 clusters, most of which contained cells predominantly from a single OD (Figure 2B). The most notable exceptions are two smaller clusters that contain cells from multiple ODs: a cluster with cells from OD2-3.2 that differentially express myo-inositol metabolism pathway genes, and a very small cluster containing only 36 cells from 5 different OD points uniquely expressing genes associated with the defective PBSX prophage (Figures 2B **and S9-S12**).

**Fig. 2.**
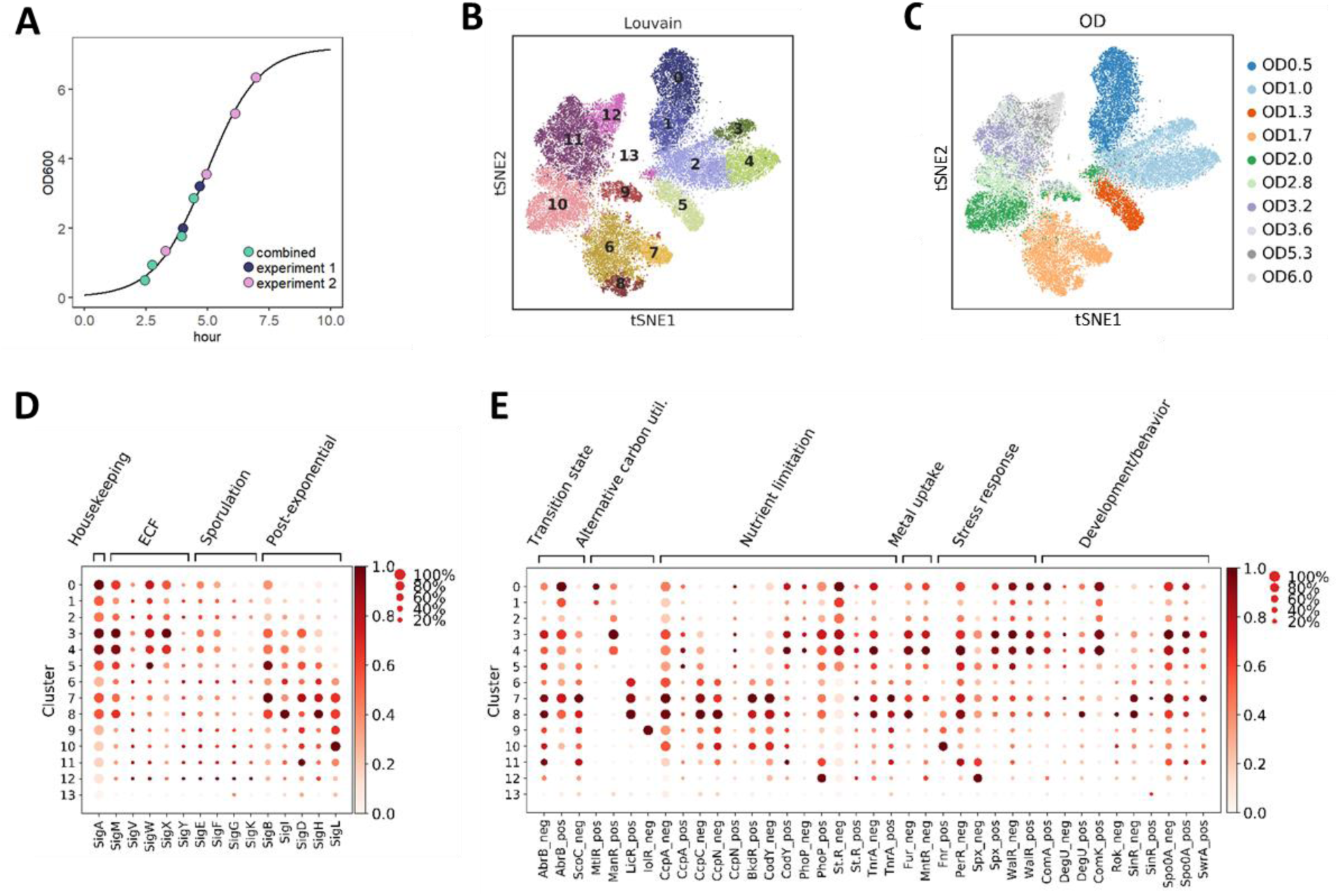
microSPLiT detects global transcriptional states during *B. subtilis* growth. **(A)** Optical density (OD) points sampled in two experiments overlayed on the growth curve from experiment 2 (see Supplementary Text). (**B**) t-SNE embedding of the combined growth curve data colored by cluster. A total of 14 clusters were found using Louvain clustering. (**C**) Inferred normalized sigma factor activity for each cluster. The size of each dot indicates the proportion of cells in the cluster in which the sigma factor is active, while the color indicates the average activity. (**D**) Inferred normalized activity of select transcriptional regulators per cluster. “Neg” indicates that activity was calculated for the genes known to be negatively regulated by this TR, and “pos” indicates the activity was calculated for the genes positively regulated by the given TR.

Alternative sigma factors are the primary regulators of the prokaryotic RNA polymerase specificity and thus directly shape transcriptional changes in response to environmental conditions. To understand whether microSPLiT could capture variation in sigma factor utilization across different growth stages, we averaged activity of genes regulated by each sigma factor, recording, for each cluster, both the percentage of cells expressing at least one gene regulated by a given sigma factor and the average intensity of gene expression (Figure 2C). We note that while averaged intensity could also be obtained from bulk RNA-seq, only single-cell methods can reveal information about cell-to-cell variation in regulon activity.

Consistent with expectations, we observe that the housekeeping σ^A^ activity is highest at OD0.5, while the activity of general stress response sigma factor σ^B^ rises as cells begin to exit from exponential phase (OD1.3-1.7) and then declines as cells approach stationary. Sporulation sigma factors σ^F^, σ^J^ and σ^K^ were induced at later ODs but in a small fraction of cells, consistent with the previously reported heterogeneous initiation of sporulation (*3*, *20*). The extracellular function (ECF) sigma factors σ^M^, σ^W^ and σ^X^ implicated in maintaining cell envelope function reached maximal activity at OD 1.0 in a large proportion of cells, consistent with reports of their basal activity in logarithmic phase in non-stressed cells (*21*). Meanwhile, the remaining ECF sigma factors σ^V^, raising defenses against lytic endoglycosidases, and σ^Y^ increased in activity towards later OD points in a small subpopulation of cells, similar to the sporulation sigma factors. σ^I^ and σ^H^ activities, regulating heat response and post-exponential behavior respectively, peak in cluster 8 which represents a subgroup of cells at OD 1.7. In contrast, σ^B^ and σ^D^ regulating general stress response and motility are most active in cluster 7, a second distinct subgroup of OD 1.7 cells. Finally, σ^L^ implicated in utilization of arginine, acetoin and fructose as well as regulation of the cold shock response peaks in cells at OD 2-2.8 represented in cluster 10, most likely due to the highly enriched acetoin utilization genes in this cluster. Additionally, we found that correlations between the sigma factor regulons largely agreed with the concept of molecular time sharing, i.e. the idea that sigma factors compete for RNA polymerases (**Figure S7**) (*22*).

To obtain an even finer-grained picture of the transcriptional programs during exponential growth and entry to stationary phase, we inferred the activity profiles of select transcriptional regulators (TR) from expression of the genes in their respective regulons (Figure 2D **and S8**) (*22*). This analysis revealed pronounced changes in regulation of carbon utilization, stress responses, metal uptake, developmental decisions and more. The main transition state regulator AbrB becomes inactive after OD1.0, indicating that the preferred carbon sources such as glucose start getting depleted. In response, cells begin to activate transcriptional programs to utilize different alternative carbon sources. Toward the intermediate growth stages, the cells start to sense carbon, nitrogen and phosphate limitation. Clusters 7 and 8 (OD 1.7) display a striking change in carbon metabolism indicated by the strong expression of genes repressed by the carbon catabolite control proteins CcpA, CcpC and CcpN. Similarly, the regulator of nitrogen assimilation TnrA becomes activated at OD1.7, while PhoP, regulating the phosphate metabolism, becomes active at three different growth stages: OD1.0, OD1.7 and later on at OD6.0. In addition, cells respond to metal deficiency, switching off the negative regulators of iron and manganese uptake Fur and MntR after OD1.0. Finally, we can observe cellular response to a variety of intrinsic and cell-envelope stresses, as well as temporal activation patterns of a battery of developmental regulators including ComA, SinI, DegU, Rok, Spo0A and others. Surprisingly, we also observe an upregulation of ComK-regulated genes in a high proportion of cells in the early ODs which is not consistent with the primary role of this transcription factor in a rare developmental state of competence. It could be explained by the fact that a large cohort of ComK-induced genes are involved in metabolism and DNA repair and can be activated by other regulators. These data show that microSPLiT not only captures known regulatory programs but also reveals intriguing heterogeneity in a wide range of these pathways.

### Central carbon metabolism changes and differential expression of tricarboxylic acid (TCA) cycle enzymes

Given the dramatic changes in regulation of carbon utilization observed in our TR analysis, we turned to a more comprehensive examination of carbon metabolism genes enriched in each cluster (**Figures S9-S12 and** Figure 3). When glucose and other preferred sugars (discussed below) are present, they are converted to pyruvate during glycolysis, which is the primary metabolic route under excess of these sugars. In these conditions promoting rapid growth, pyruvate is then converted to lactate, acetate, acetoin, and other by-products through fermentation. Upon depletion of preferred sugars, cells redirect the fermentation by-products to be metabolized in the TCA cycle generating additional adenosine triphosphate (ATP) and carbon dioxide.

**Fig. 3.**
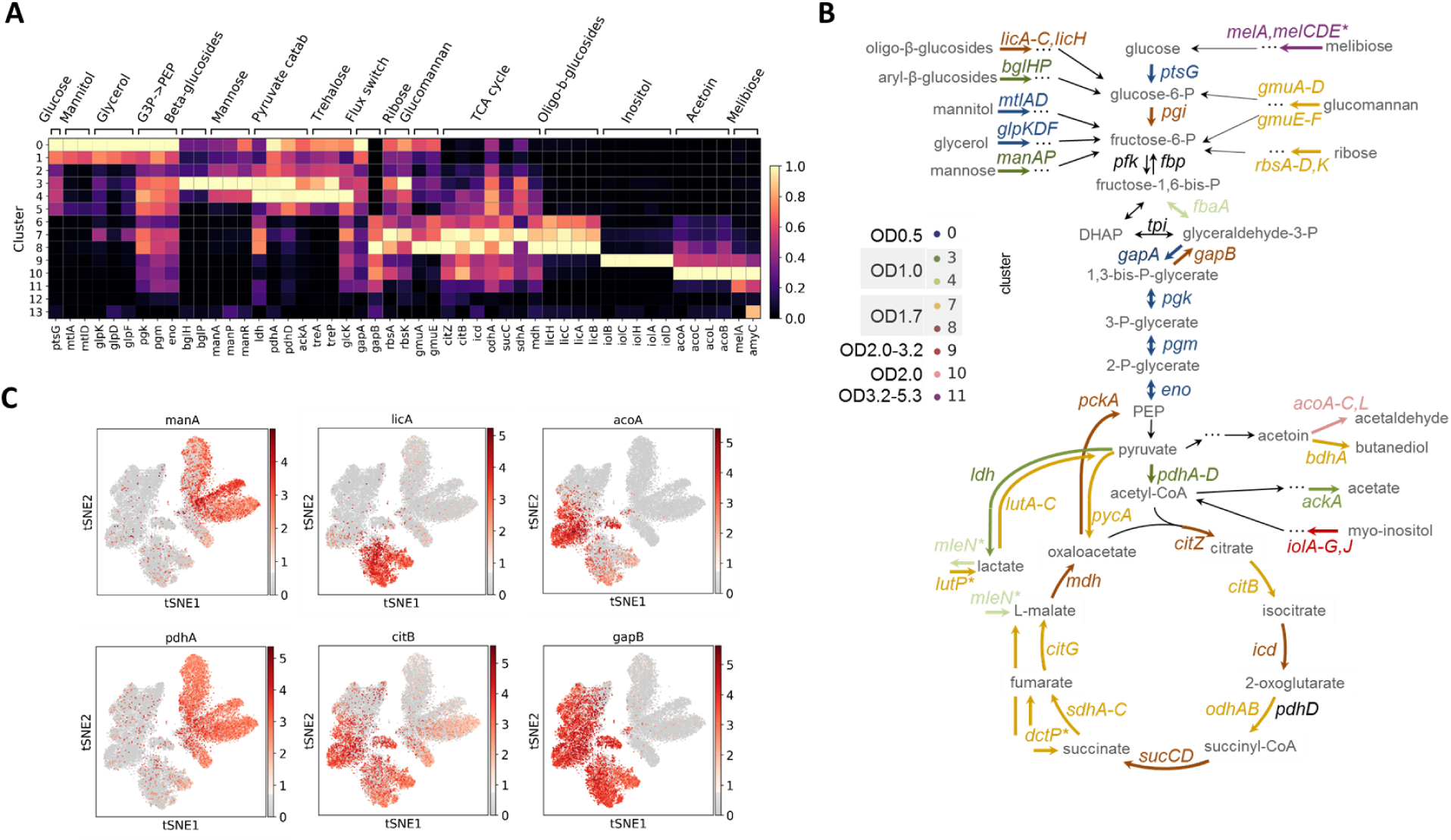
Central carbon metabolism changes and alternative carbon sources utilization during *B. subtilis* growth. **(A)** Normalized expression of genes from select metabolic pathways and central carbon metabolism shown per cluster. Gene expression clearly shows distinct carbon utilization programs associated with different clusters and growth states. (**B**) Expression of select genes from (A) overlaid on the t-SNE plot to illustrate the differential patterns of activation. (**C**) Schematic of the central carbon metabolism pathway showing alternative carbon sources, metabolic products and genes in the pathway. The genes are color-coded according to the cluster they are highest expressed in.

In the exponential phase (Clusters 0 and 1, OD0.5) we find high expression of *ptsG*, a glucose permease which transports and phosphorylates glucose. The enzymes in the *gapA* operon constituting the metabolic pathway from glyceraldehyde-3P to phosphoenolpyruvate (PEP): *gapA, pgk, pgm* and *eno* (*23*) were upregulated in this cluster (Figure 3A,B).

Next, in clusters 3 and 4 (OD1.0) we observe transcriptional patterns suggesting an increase in flux from pyruvate either being converted to lactate by *ldh* (*24*) which is then exported via a malate antiporter *mleN* (*25*) or converted to acetate via intermediates by *pdhAD* (*26*) and *ackA* (*27*) (Figures 3A,B). These observations are consistent with previous reports of transient medium acidification via acetate production during rapid fermentative *B. subtilis* growth (*28*).

At OD1.7 cells appear to undergo a dramatic transition from glycolysis to gluconeogenesis with multiple genes from the gluconeogenetic pathway activated in clusters 7 and 8. There are two glyceraldehyde-3-phosphate dehydrogenases in *B. subtilis*: GapA and GapB, mediating the flux of carbon either from glucose to the TCA cycle or vice versa (*29*). The glucose and intermediates generated by the gluconeogenetic pathway under conditions of glucose limitation are then used for synthesis of necessary structural constituents. We observe the switch from GapA to GapB expression in clusters 7 and 8 along with an upregulation of most of the TCA cycle enzymes. We also find a different pattern of pyruvate production and utilization, together with catabolism of acetoin, another fermentation product, and additional nutrient fluxes into the TCA cycle (**Supplementary Text and** Figures 3A-C, **S9-S12**).

Interestingly, the TCA cycle enzymes which are collectively enriched in the entire OD1.7 sample display differential transcriptional abundances between clusters 7 and 8 (Figures 3A, B). For instance, cells in cluster 7 express more aconitase (*citB*), 2-oxoglutarate dehydrogenase (*odhAB*), and succinate dehydrogenase (*sdhABC*), while cells from cluster 8 display higher expression of genes in the *citZ-icd-mdh* operon and *sucCD* (succinyl-CoA synthetase). CitB was reported to destabilize the *citZ-icd-mdh* mRNA upon citrate accumulation or iron limitation (*30*). Thus, differential transcript abundance of the TCA cycle enzymes could reflect the regulatory interactions between the respective genes.

### Transient activation of alternative carbon utilization pathways in distinct subpopulations

Complementing changes in the core carbon metabolism, we observe expression of pathways responsible for uptake and utilization of a variety of different carbon sources. As the preferred sources of carbon are depleted, the major repressor of alternative carbon utilization pathways CcpA becomes inactive, permitting the cells to catabolize alternative carbon sources including a variety of carbohydrates (*31*) (Figure 3). We find that the activation and suppression of these pathways happen in varying proportions of the cells in each OD sample and appear to follow a temporal order (Figures 3 **and S9-S13**). The preferred carbon sources for *B. subtilis* are glucose along with other Group A sugars (sucrose, fructose, and mannitol) and malate (*32*, *33*). While the latter is directly metabolized in the TCA cycle, the former sugars are converted to one of the glucose metabolic intermediates. At OD0.5, in addition to glucose-specific *ptsG*, we observe increased expression of *mtlA-D* and *glpK,D,F* responsible for utilization of mannitol and glycerol, respectively. Cells in clusters 3 and 4 (OD1.0) activate catabolism of mannose and aryl-β-glucosides (*manA,P* and *bglH,P*). In cluster 7 (OD1.7) we observe the upregulation of genes for utilization of glucomannan (*gmuA-F*) and the ribose transporter (*rbsA-D*), while the gene for utilization of ribose *rbsK*, curiously, is upregulated much earlier in cluster 3. Finally, at even later ODs three additional alternative carbon source utilization programs switch on. Cluster 9 comprising cells from ODs 1.7, 2.0, 2.8, and 3.2 is defined by the expression of genes implicated in the most common stereoisomer of inositol, myo-inositol catabolism (*iolABCDEFGHIJ*, further “*iolAJ*”), while cluster 10 (OD2.0) is enriched for genes responsible for utilization of acetoin (*acoABCL*). Finally, cluster 11, representing a range of ODs from 3.2 to 5.3, differentially expresses genes for melibiose utilization (*melA, melCDE*).

### Heterogeneous activation of myo-inositol catabolism pathway at intermediate growth stages

Inositol is an abundant resource in soil, and *B. subtilis* is able to subsist on inositol as its sole carbon source (*34*). While LB medium is not typically expected to contain myo-inositol (further “inositol”), heterogeneous inositol utilization pathway activation is observed in a small (3-15%) subpopulation in both of our independent LB growth experiments (cluster 9, see Figure 3A, Figure 4A-B and **Figure S14**). It has been shown that the inositol catabolism intermediate, 2-deoxy-5-keto-D-gluconic acid 6-phosphate (DKGP), is responsible for the pathway induction (*34*). We hypothesize that the trace amounts of inositol may be present in the LB medium, potentially from the yeast extract since yeast is capable of inositol production as a precursor to the essential membrane component, phosphatidylinositol.

**Fig. 4.**
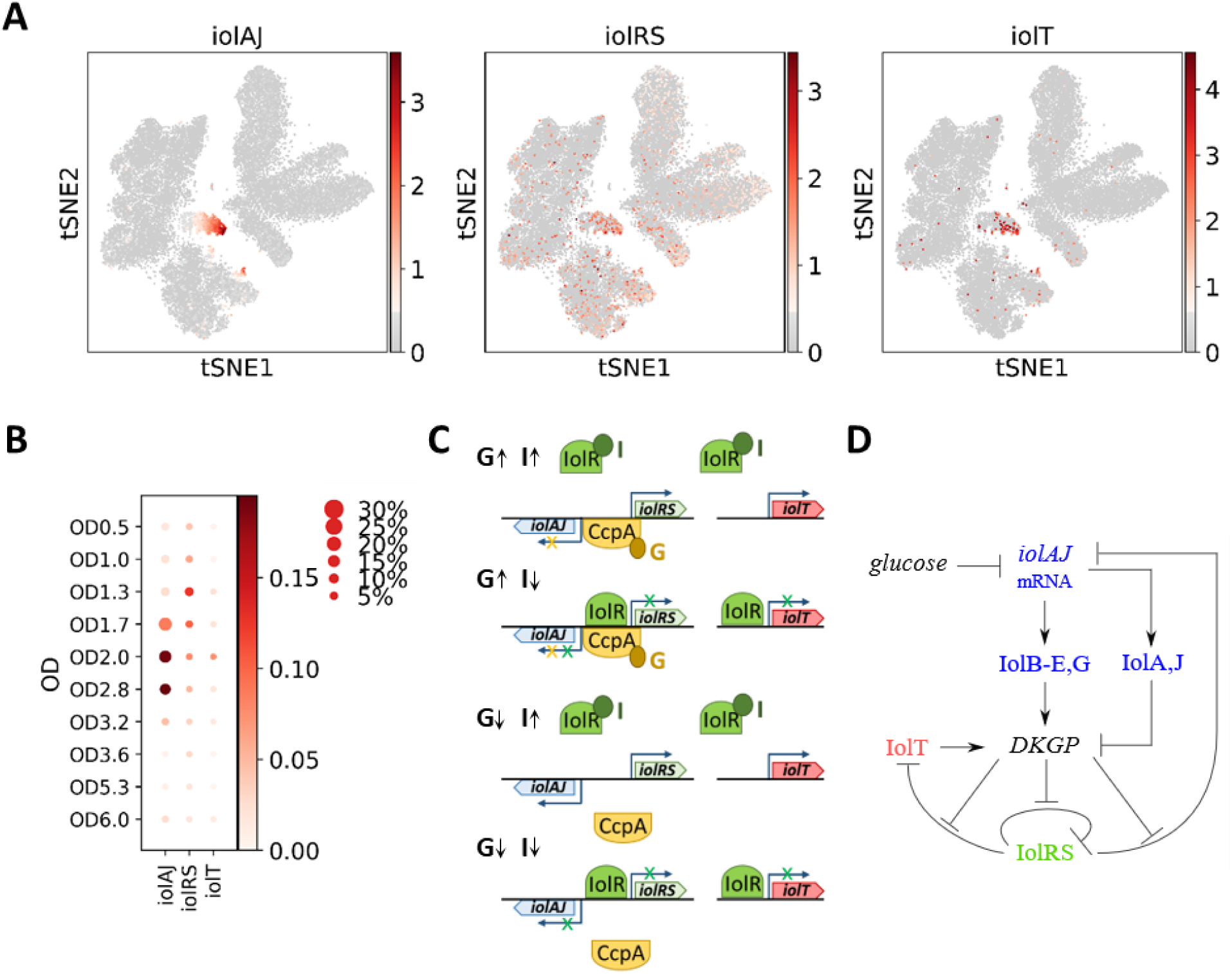
Myo-inositol utilization pathway genes are expressed heterogeneously at intermediate growth stages. **(A)** Expression of each of the three inositol utilization operons, averaged across all genes in a given operon, and overlaid on the t-SNE plot. (**B**) Activities of the three inositol utilization operons across ODs. The size of each dot indicates the proportion of cells in each OD sample expressing any of the genes in the selected operon, while the color shows the average expression of the genes in a given operon. (**C**) Schematic of the inositol utilization operons structure along with the main regulators in different metabolic states. When both glucose (G) and inositol (I) are present in the cell, the IolR repression of all three operons is relieved, but CcpA still represses the *iolAJ* operon preventing it from metabolizing inositol (we show the signaling intermediates bound to the repressors also as G and I for simplicity). As glucose levels drop, CcpA no longer inhibits transcription of *iolAJ*, and if inositol is present, it relieves the repression of *iolRS* and *iolT*, inducing the expression of the entire metabolic network. Meanwhile, in the absence of inositol, all of the operons remain repressed regardless of the glucose levels. (**D**) A simplified network diagram representing the interactions and regulations within the inositol utilization pathway.

There are three operons involved in inositol utilization, *iolT* (main transporter), *iolRS* (the first gene is a repressor and the second is a likely dehydrogenase), and *iolAJ* (metabolic enzymes), with *iolC* producing and *iolJ* cleaving the pathway-activating DKGP intermediate. *iolRS* and *iolAJ* are normally transcribed by σ^A^ through divergent transcription (*35*). In the absence of the inducer (I), IolR supresses transcription of all three operons (*36*). In addition, CcpA represses the *iolAJ* operon in the presence of glucose (*37*) (Figure 4C). Interestingly, the pathway suppressor *iolR* gene is more broadly expressed both outside and inside of cluster 9 (Figure 4A-B and **Figure S12**).

Next, we asked whether our data could be explained by the underlying gene regulatory network architecture (Figure 4D). Within the complex system of interactions, we observe two topological features capable of amplifying small molecular variations. An *iolAJ*-mediated incoherent feed-forward loop can generate a pulse of DKGP, while IolT can generate a positive-feedback loop by transporting inositol into the cell. These features of the network can explain the difference in clusters, as probed with a qualitative model (**Figure S15**). Due to the small proportion of cells expressing the main catabolic operon *iolAJ*, this particular metabolic behavior could only be reliably detected on a single-cell level.

### Motility, antimicrobials production, stress response, and metal ion import

*B. subtilis* is known to exhibit a variety of behaviors to enhance survival in adverse conditions including, but not limited to, production of degradative enzymes and antimicrobials, secretion and uptake of siderophores, three types of motility (swimming, swarming, and sliding), natural competence, and sporulation (*38*). Many if not most of these behaviors are not displayed by all cells at the same time (*39*), thus providing good targets for single-cell interrogation.

Bacteria universally produce peptide and small molecule antimicrobials that are meant to target both closely and distantly related organisms (Figure 5A) (*40*). We observe the expression of endoA toxin/antitoxin (*ndoA/I*) and bacilysin (*bacA-G*) more broadly prior to OD2.0 (Figure 5A **and S16**). Subtilosin (*sboA,X*-*albA-F*), bacillaene (*pksC-R*), and plipastatin (*ppsA-E*) production, on the other hand, are detected in a greater proportion of cells after OD2.0. All three of these peptides are broad spectrum antimicrobials (*41*–*43*) which suggests that under nutrient limitation, *B. subtilis* becomes competitive to preserve its access to the available resources, even in the absence of other species. Additionally, we also see a rise in spore killing factor (SKF) and spore delay protein (SDP) in the last three ODs (**Figure S17**).

**Fig. 5.**
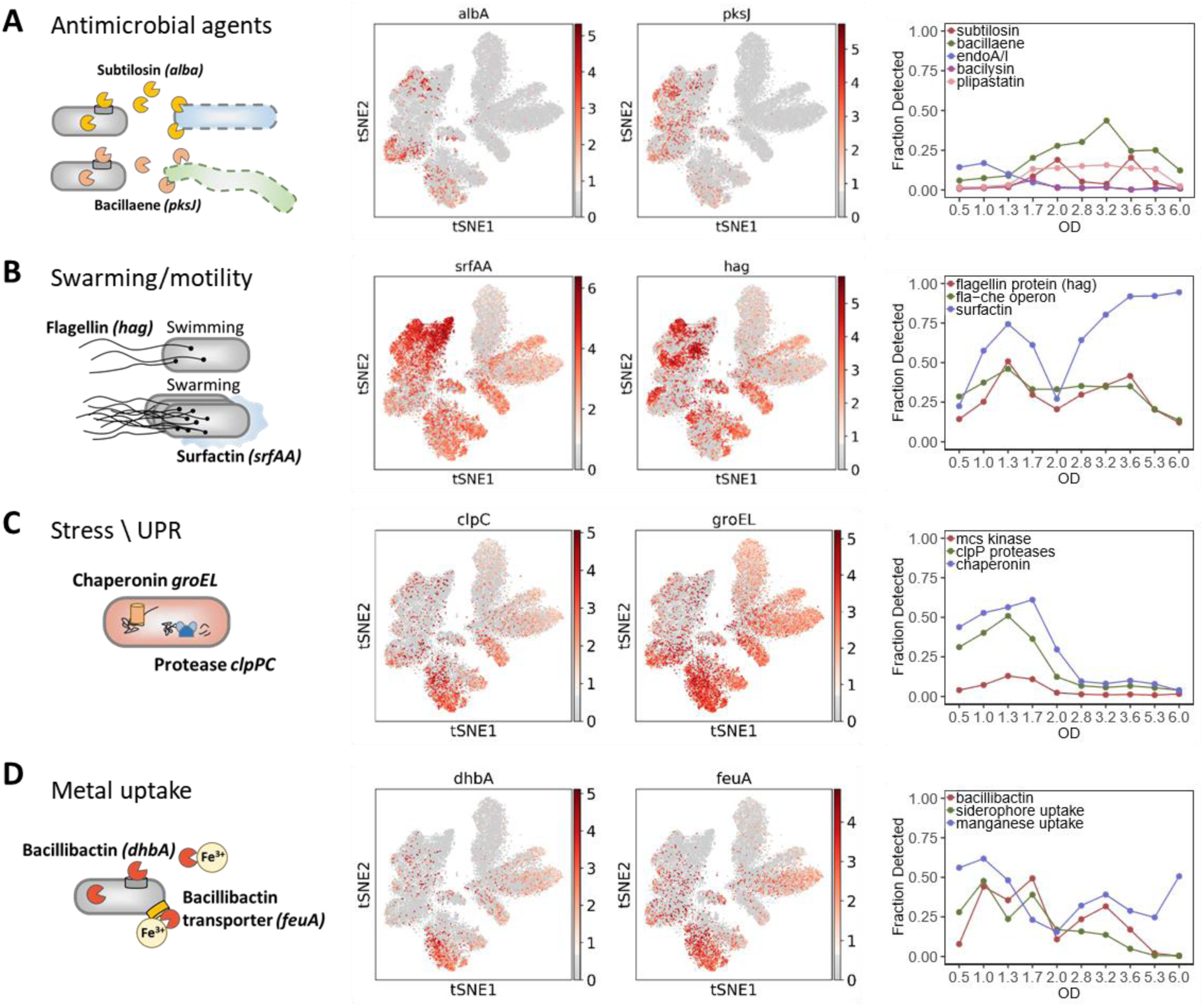
Intrinsic stress responses and developmental gene expression. **(A)** Antimicrobial agents and endoA toxin-antitoxin secretion. Subtilosin and bacillaene action diagram. Overlay of representative subtilosin (*albA*) and bacillaene (*pksJ*) production pathway gene expression on the t-SNE. Fraction of cells expressing the indicated antimicrobials and endotoxin-antitoxin as a function of OD. **(B)** Swarming and motility genes expression. Flagellin and surfactin action diagram. Overlay of surfactin (*srfAA*) and flagellin (*hag*) gene expression on the t-SNE. Fraction of cells expressing the indicated motility operons as a function of OD. **(C)** Intrinsic stress and unfolded protein response (UPR) genes expression. GroEL chaperonin and ClpCP protease action diagram. Overlay of *clpC* and *groEL* gene expression on the t-SNE. Fraction of cells expressing the indicated genes as a function of OD. **(D)** Metal (iron and manganese) uptake genes expression. Bacillibactin siderophore and transporter action diagram. Overlay of bacillibactin (*dhbA*) and siderophore transporter (*feuA*) production pathway gene expression on the t-SNE. Fraction of cells expressing the indicated metal uptake genes as a function of OD. The genes used for fractional plots are listed in Supplementary Table S2.

*B. subtilis* has two morphological phenotypes during active growth: filamentous, sessile cells and smaller motile cells (*44*, *45*). In liquid media, the bacteria primarily swim which requires the expression of the flagellin protein (*hag*) and the *fla-che* operon (Figures 5B). We detected expression of these genes in 25 to 50% of cells in every cluster from early to intermediate growth stages before the fraction noticeably declines at late ODs, consistent with reports of heterogeneity of the motility operon expression at mid-exponential phase (*42*). The *B. subtilis* population is expected to be similarly differentiated into surfactin-producing and extracellular matrix-producing bacteria as cell density increases (*46*). We see the fraction of cells producing *srfA-D* genes gradually increase, slightly dipping at OD 1.7 and 2.0, and finally reaching almost 100% detection at OD 6.0, consistent with the PY79 strain having defective matrix production genes that cannot negatively regulate *srfA-D* expression (*47*).

Cellular stress response pathways exemplified by GroEL chaperonin and ClpC proteases peak at OD1.7, the same time as the cells switch from glycolysis to gluconeogenesis (Figure 5C). The ClpP associated proteases (*clpP,C,X,E*), McsA and McsB kinases (*mcsA,B*), and chaperonins (*groEL,ES*) are all involved in the unfolded protein response (*48*). A transient increase in the regulatory sigma factor, σ^B^, inducing expression of these genes, has been reported during normal exponential growth and attributed to intrinsic cellular stresses (*49*). *clpP* mutants have been linked to slower glucose consumption and overproduction of TCA cycle metabolites during rapid fermentative growth, which suggests their participation in glycolysis and may explain their highest expression during the early growth stages (*50*).

*B. subtilis* needs both manganese and iron to grow (*51*), and the two ions often antagonize each other’s regulators (*52*). Upon iron limitation, *B. subtilis* produces and secretes siderophores including bacillibactin (*dhbA*) together with the ABC transporter (*feuA*) (Figure 5D) (*53*, *54*). Both genes show increased expression at OD1.7, similar to the stress response proteins above. Manganese, on the other hand, is transported into the cells via a separate ABC transporter (*mntA-D*) and a proton symporter (*mntH*) (*52*). It is required for the successful transition into stress related states such as biofilm formation and sporulation (*55*), which likely explains the increase in manganese transport related genes detected at later ODs.

### microSPLiT quantifies a rare stress response

Cluster 13 (36 cells, or 0.142% of total cells, representing ODs between 0.5 and 2.8) contains a rare subpopulation of cells expressing PBSX prophage genes (Figure 6A). The PBSX element is a defective prophage that is non-infectious but upon induction causes the release of phage-like particles (*56*) containing 13 kb of random fragmented chromosomal DNA (*57*, *58*). Prophage gene expression is induced by DNA damage mediated by the SOS response (*56*, *59*)(Figure 6B). We identified eleven host genes with known or putative functions expressed in the PBSX prophage cluster (Figure 6C). Five of these genes have previously been shown to be induced only in PBSX-harboring strains of B. S. after DNA damage (*57*). The rest, including a chemoreceptor (*mcpC*), an ATP-binding cassette transporter (*liaL*), a cell wall binding protein (*ykuG*), an ammonium transporter (*amtB*), a sucrose-6-phospate hydrolase (*sacA*), and a regulatory protein of homologous recombination (*recX*), have not previously been linked to prophage induction (**Supplementary Text**). These genes could have been induced by the damage sustained by the cells which also caused PBSX prophage activation. Alternatively, these genes could have been activated in response to the chromosomal fragmentation and membrane vesicle formation caused by the prophage induction.

**Fig. 6.**
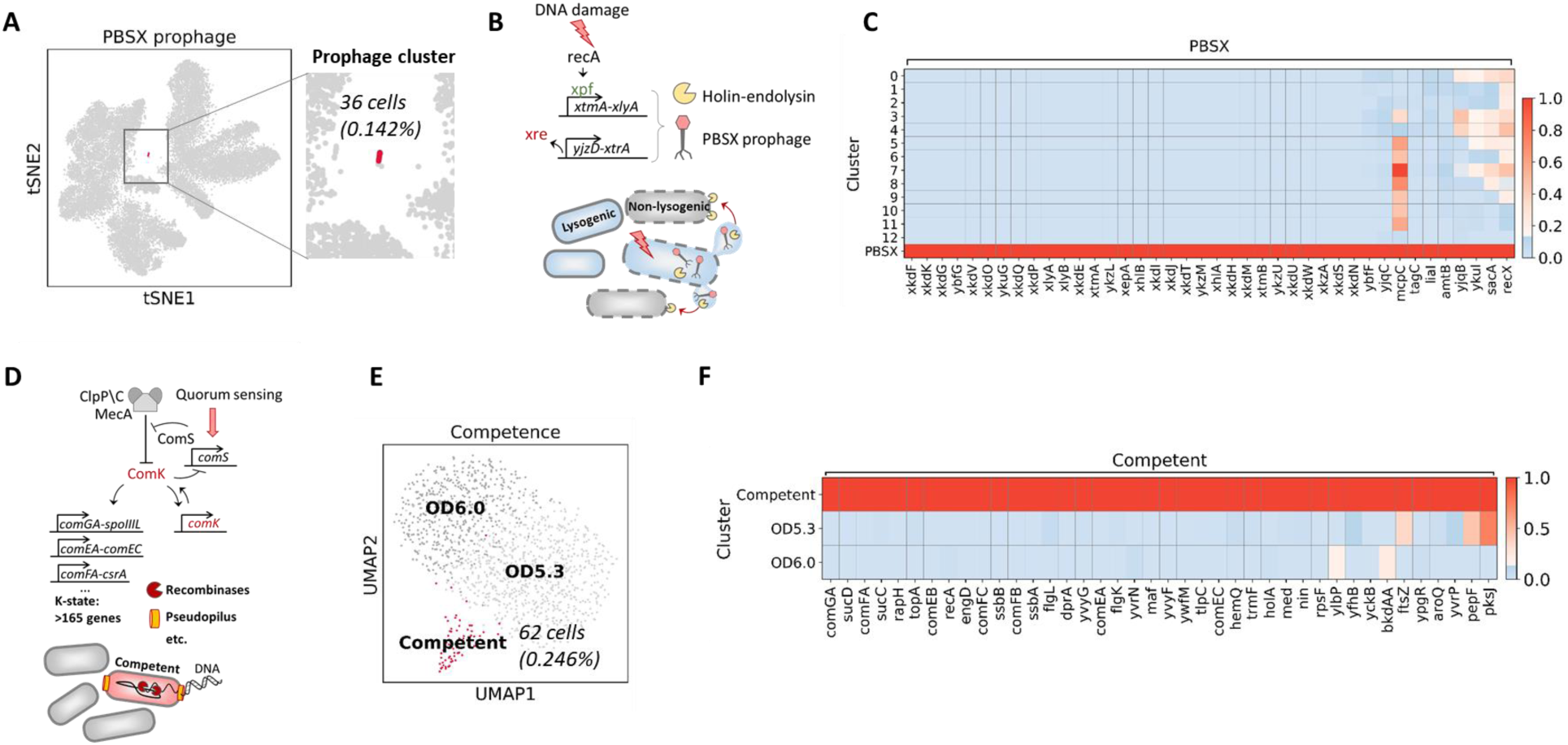
Rare developmental states induced by cellular stress. (**A**) PBSX prophage induction and killing of the non-lysogenic cells. Upon DNA damage, two large prophage operons are induced and non-infectious phage particles along with endolysins are released. The latter degrade cell walls of other cells containing heterologous prophage elements. (**B**) PBSX prophage cluster (36 cells) shown on the t-SNE plot. (**C**) Genes enriched in the PBSX prophage cluster, including both prophage and host genes. (**D**) Overview of competence development. Activated by stress and quorum-sensing, ComS binds to and redirects MecA complex from degrading master competence regulator ComK. Amplified via positive autoregulation, ComK induces >165 genes involved in extracellular DNA uptake and cellular physiology. (**E**) UMAP embedding of the subclustered OD5.3 and 6.0 samples, showing the competence cluster (62 cells). (**F**) Genes enriched in the discovered competence cluster relative to the rest of the cells in OD5.3 and 6.0 samples.

### microSPLiT detects a rare stochastically induced developmental state

Under stress or nutrient limitation, a small fraction (2-5%) of *B. subtilis* cells undergoes stochastic transient differentiation into a state of natural competence, characterized by the ability to uptake extracellular DNA and integrate it into the chromosome (Figure 6D) (*60*). The master transcriptional regulator of competence ComK is activated via a positive feedback loop, inducing expression of a suite of >165 genes involved in a variety of cellular processes in addition to DNA uptake (*61*). Competence is expected to naturally occur under nutrient limitation. We thus separately subclustered the last two OD points (OD5.3 and 6.0). UMAP embedding revealed a small cluster (62 cells, or 4.6% of cells at OD 5.3 and 6) expressing a distinct transcriptional signature of the competent state, or K-state (Figure 6E-F). The most enriched gene was *comGA*, as expected from prior transcriptomic data, followed by the succinyl-CoA synthetase (*sucCD*) operon which has been shown to be induced in competent cells (*28*, *61*, *62*). We also see strong enrichment of other genes encoding the DNA uptake machinery: *comF* and *comE* operons, as well as the response regulator RapH which represses sporulation development in competent cells by dephosphorylating the phosphorelay component Spo0F (*63*). As the cells in the competent state arrest replication to integrate the incoming DNA into the chromosome, we observe the upregulation of genes necessary for processing of internalized ssDNA such as DNA recombination and repair protein RecA along with single-strand DNA binding proteins SsbA and SsbB consistent with prior reports (*61*). We also observe enrichment of several genes related to DNA processing such as *topA* encoding topoisomerase A and *holA*, delta subunit of DNA polymerase III which is a part of the replisome. These genes are not in the annotated ComK regulon but were tentatively identified in the microarray data comparing gene expression between *mecA* strain in which essentially all cells express ComK and a double mutant *mecA comK* strain (*61*). This coordinated upregulation of *topA* and *holA* is consistent with the model of RecA binding to the SsbA\SsbB coated ssDNA and forming a complex with the replisome during competence (*64*). In addition, we found four genes not previously linked to the competent state: *ywfM* (unknown), *hemQ* (coproheme decarboxylase), *tlpC* (an orphan membrane-bound chemotaxis receptor), and *trmF*, a folate- and FAD-dependent tRNA methyltransferase (**Supplementary Text**).

## Discussion

We applied microSPLiT to *B. subtilis* cells growing in liquid rich medium, conditions which are not usually associated with abundant cellular heterogeneity. Nevertheless, we found a large variety of subpopulations displaying differential gene expression of select metabolic, stress response or developmental pathways. In particular, we identified a metabolic pathway, myo-inositol catabolism, which was strikingly activated only in a fraction of cells from several later OD points in a distinct temporal fashion. We expect microSPLiT to have even more utility in identifying heterogeneous cell states in an environment more conducive to bacterial differentiation, such as in multi-species biofilms and natural microbiota.

We were able to detect subpopulations of cells as rare as 0.142%, pointing to microSPLiT’s potential to uncover physiologically relevant rare cell states, such as persistence, that are hard to study by bulk or low-throughput methods. For many such states, the regulators are not well known and consequently, the reporters or mutants producing the desired state at a higher frequency cannot be engineered. Even for states that are better understood and can be artificially induced, such as prophage induction by UV irradiation, microSPLiT is useful since the transcriptional signatures it produces are state-specific and free of artifacts introduced by the perturbation.

In order to use microSPLiT on complex natural communities, the protocol will likely need to be further optimized, particularly the permeabilization and mRNA enrichment steps. Since cell wall and membrane composition vary significantly among bacteria, any permeabilization treatment is likely to work better for some species than others. However, alternate treatment for different subsamples may still provide optimal results. In addition, we experienced lower mRNA counts from bacteria in stationary phase as opposed to logarithmic growth phase (**Figure S18**), consistent with slower growth rate and smaller cell size at this stage. Although the resulting data were still sufficient to reliably identify rare cell states such as the K-state, further improvement of the protocol will be instrumental to increase sensitivity for applications to slower dividing bacteria or challenging environmental conditions. Still, we expect microSPLiT to provide an exciting new dimension to studies of bacterial gene expression heterogeneity and community behavior facilitated by the method’s potential scalability to millions of bacterial cells and single-cell resolution without need for constructing reporters.

## Supporting information

Auxiliary Table S3

Auxiliary Table S4

## Acknowledgments

This work was supported by NIH grants NIH R01HG009136 and R01HG009892 to G.S. and NIH R01DK104908-01 to W.D. All relevant sequencing files will be deposited to Sequence Read Archive (SRA). Processed data will be submitted to GEO. A.K., L.B. W.D. and G.S. have filed a patent application for microSPLiT. C.R., A.B.R and G.S. are co-founders of Split Biosciences, a single-cell RNA sequencing company.

## Supplementary Materials

Materials and Methods

Figures S1-S18

Tables S1 and S2

Tables (Auxiliary Excel Files) S3 and S4

Supplementary References (*1-29*)

## Notes

#### Summary of Updates

Figure 6 revised; supplemental files updated; supplemental files added.

