## Supplementary Materials for "Microbial single-cell RNA sequencing by split-pool barcoding"

**This PDF file includes:**

Materials and Methods
Supplementary Text
Figures S1-S18
Tables S1-S2
Supplemental References

**Other Supplementary Materials for this manuscript includes the following:**

Tables (Auxiliary Excel Files) S3 and S4

### Materials and Methods

### Experimental Methods

#### Oligonucleotides

The sequences and modifications for all oligonucleotides used can be found in Auxilary Table S3.

#### Bacterial Culture

*Escherichia coli* MW1255, a derivative of MG1655 (E.C.), and *Bacillus subtilis* PY79 (B.S.) overnight cultures were inoculated into fresh LB medium at 1:1000 or 1:250, respectively, and grown at 37C with shaking (200rpm). For the heat shock experiment, upon reaching the $OD_{600}$ = 0.5 half of each culture was transferred to the separate 37C incubator where the temperature was increased to 47C, and kept there with shaking (200rpm) for 8 minutes from the time the temperature stabilized. Both control and heat-treated samples then were immediately centrifuged at 4C, 5000 rcf for 5 minutes, and resuspended in cold formaldehyde. For the B.S. growth curve, samples were centrifuged at room temperature, 10000 rpm for 1 minute before fixation.

#### Fixation and permeabilization

For the steps below, centrifugation was performed at 4C, 5000 rcf for 5 minutes. Following centrifugation, the bacterial pellet was resuspended in 1mL of fresh, cold, 4% formaldehyde solution (in 1x PBS) and incubated at 4C overnight. The next morning, cells were centrifuged and resuspended in 1ml cold 100mM Tris-HCL+RI ('RI' indicates that SUPERase-In RNase Inhibitor, ThermoFisher, was added to a final concentration of 0.1 U/uL). Cells were centrifuged and resuspended in 250ul of 0.04% Tween-20 in 1x PBS, then permeabilized for 3 min on ice. We then added 1ml of cold PBS+RI, centrifuged the cells and resuspended in 200ul lysozyme mix per sample on ice as follows: 0.1M Tris-HCL pH7, 0.05M EDTA, 2.5mg/ml lysozyme, 0.25U/ml SUPERase-In. We incubated the samples at 37C in the thermocycler for 15 minutes as we found that precise timing of lysozyme incubation is critical to maintaining cell integrity at later stages of the protocol. Following the cell wall digestion step, we immediately added 1ml of cold PBS+RI, centrifuged the cells and counted the cells stained with SYTO9 using Accuri C6 flow cytometer. For the species-mixing experiment, we mixed the E.C. and B.S. cells at equal proportions and took 0.6M cells for each of heat shocked and control samples. For the B.S. growth curve experiments, we took 0.25M cells for each OD sample.

#### In-cell Polyadenylation

In order to enrich for mRNA capture, we performed *in situ* polyadenylation with *E. coli* Poly(A) Polymerase I (PAP) from NEB. For 0.25M cell samples, the reaction proceeded in 50µl volume, and in 100µl volume for the 0.6M cell samples. For each 0.6M cells bacterial pellet, we added 66µl of water, 4µl of SUPERase-In, 10µl of 10x PAP Buffer, 10µl of 10mM ATP, and 10µl of PAP. The reaction mixture was incubated at 37C for 30min, then centrifuged upon addition of 1ml cold PBS+RI. We also added 1µL of 10% Tween-20 in order to make the cells easier to pellet. The cells then were resuspended in 0.5ml of cold PBS+RI.

#### Reducing aggregate formation

Two steps were crucial to break down cell aggregates and reduce the doublet rate: first, we vortexed and double-filtered the bacterial cells prior to reverse transcription; and second, we performed sonication with double filtration right after the reverse transcription.

Following the polyadenylation step, cells in 500ul of PBS-RI were vortexed for 1 minute on the highest setting, filtered through 10µM pluriStrainer (pluriSelect) by pipetting through the membrane, then filtered again through 1µM pluriStrainer with gentle suction, and finally, right before adding to the reverse transcription wells, vortexed again on the highest setting for 1 minute.

Following the reverse transcription step and after resuspension in 2 mL of cold PBS+RI, cells were vortexed for 1 minute on the highest setting, filtered through 10µM and 1µM pluriStrainer as above, and briefly sonicated at 10% power for 5s on ice for 1 pulse (Sharpertek Ultrasonic Cell Crusher) followed by immediate distribution to the ligation plate. We found that the sonication step can be replaced by a second vigorous vortexing step, with the roughly 2-fold increase in resulting detected doublet rate (0.7% to 1.3%).

<u>In-cell Reverse Transcription</u>
Like in SPLiT-seq, the first round of barcoding occurs through an *in situ* reverse transcription (RT) reaction. Cells are split into up to 48 wells, each containing barcoded well-specific reverse transcription primers. We used both random hexamer and anchored poly$(dT)_{15}$ barcoded RT primers in each well at the ratio of 1:2 (2.5µM ramdom hexamer + 5µM poly$(dT)_{15}$). In addition to primers, each RT well had a mix of 1X RT Buffer, 0.25U/µL RNase Inhibitor (Enzymatics), 0.25U/µL SUPERase-In RNase Inhibitor, 500µM dNTPs each (ThermoFisher), 7.5% of PEG8000, 20U/µL of Maxima H Minus Reverse Transcriptase (ThermoFisher). We pipetted 4 µL of cells at about 1M cells per mL in PBS-RI into every well, in the total resulting RT reaction volume of 20ul. The plate was incubated in a thermocycler for 10 min at 23˚C followed by 50˚C for 50 min. RT reactions were pooled back together and after adding 9.6 µL of 10% Triton X-100, cells were centrifuged for 5 min at 3000g at 4˚C in a swinging bucket rotor centrifuge. The supernatant was removed and cells were resuspended in 2 mL of cold 1X PBS-RI. The cells then underwent two rounds of filtration and sonication as described above.

<u>Preparing Oligonucleotides for Ligations</u>
The oligonucleotide plates for the second and third barcoding round were prepared as previously described (*1*).

<u>In-cell Ligations</u>
We prepared a 2.04 mL ligation mix containing 727.5 µL of RNase-free water, 500 µL 10X T4 Ligase buffer (NEB), 20 µL T4 DNA Ligase (2000 U/µL, NEB), 30 µL RNase inhibitor (40 U/µL, Enzymatics), 12.5 µL SuperaseIn RNase Inhibitor (20 U/µL), and 750 µL of 50% PEG8000. This ligation mix and the 2 mL of sonicated and filtered cells in 1X PBS were added to a basin and mixed thoroughly to make a total of 4.04 mL.

The ligation steps were performed as in the SPLiT-seq protocol (*1*), except we did vigorous vortexing combined with the double filtration technique as described above where the protocol called for a filtration step.

<u>Lysis and Sublibrary Generation</u>
After the third round of barcoding, we performed a final vigorous vortexing and double filtration step as described above. Then 70 µL of 10% Triton-X100 was added to the cell solution before

spinning it down for 5 min at 3000G and 4˚C. We carefully aspirated the supernatant, leaving about 30 µL to avoid removing the pellet. We then resuspended the cells in 4 mL of wash buffer (4 mL of 1X PBS, 40 µL of 10% Triton X-100 and 10 µL of SUPERase-In RNase Inhibitor) and spun down for 5 min at 3000G and 4˚C. We then aspirated the supernatant and resuspended in 50 µL of PBS+RI. After counting cells, we aliquoted them into sublibraries (in 1.7 mL tubes). After adding the desired number of cells to each sublibrary, we brought the volume of each to 50 µL by adding 1x PBS and froze the cells at -80C overnight. Next morning, we flash-thawed the cells and added 50 µL of 2X lysis buffer (20 mM Tris (pH 8.0), 400 mM NaCl, 100 mM EDTA (pH 8.0), and 4.4% SDS) and 10 µL of proteinase K solution (20mg/mL). We incubated cells at 55˚C for 2 hours with shaking at 200 rpm to lyse the cells and reverse the formaldehyde crosslinks.

Purification of cDNA
cDNA purification and bonding to streptavidin beads was performed according to the SPLiT-seq protocol (*1*).

Template Switch
Streptavidin beads with bound cDNA molecules were resuspended in a solution containing 99 µl nuclease-free water, 44 µL of 5X Maxima RT buffer (ThermoFisher), 33 µL of 50% PEG8000 solution, 22 µL of 10 mM dNTPs each (ThermoFisher), 5.5 µL of RNase Inhibitor (Enzymatics), 11 µL of Maxima H Minus Reverse Transcriptase (ThermoFisher), and 5.5 µL of 100uM of a template switch primer (BC_0127). The template switch primer contains two ribonucleic guanines followed by a locked nucleic acid guanine at the end of the primer (Exiqon). The beads were incubated at room temperature for 30 minutes and then at 42°C for 90 minutes with gentle shaking.

PCR
The on-beads PCR followed by the qPCR to amplify the product were performed according to the SPLiT-seq protocol (*1*).

Illumina Sequencing
Libraries were sequenced on MiSeq or NextSeq systems (Illumina) using 150 nucleotide (nt) kits and paired-end sequencing. Read 1 (74 nt) covered the transcript sequences. Read 2 (86 nt) covered the UMI and barcode combinations. The index read (6 nt), serving as the fourth barcode, covered the sublibrary indices introduced after fragmentation.

Fragmentation
Following cDNA amplification, molecules were fragmented with a subsequent adaptor ligation step using a protocol modified from Enzymatics. Briefly, 110 ng of amplified cDNA was placed into a 50 uL reaction containing 5 uL of 10x Fragmentation Buffer (Enzymatics) and 10 uL of 5X WGS Fragmentation Mix (Enzymatics). Samples were placed into a thermocycler with the following steps: 32C for 10 min, 65C for 30 min, 4C hold. Following fragmentation, a double sided SPRI bead size selection with bounds of 0.6x – 0.8x was performed, with the final elution step using a volume of 50 uL. This 50 uL of eluant was placed into a 100 uL reaction containing 20 uL of 5X Rapid Ligation Buffer (Enzymatics), 10 uL of WGS Ligase (Enzymatics), and 2.5 uL of a pre-annealed adaptor duplex consisting of BC_243 and BC_244 at a concentration of 100 uM in 50uM NaCl. This adaptor ligation mix was incubated at 20C for 15 minutes. This was followed by a 0.8x SPRI size selection with final elution in 20 uL. Next, 18.5 uL of eluant was placed into a 50 uL qPCR reaction containing 25 uL of 2X Kapa HiFi Master Mix, 2.5 uL of 20X Evagreen, 2 uL of BC_0027 (10 uM), and 2 uL of BC_0076-BC_0083 (10 uM). This PCR mix

was placed in a thermocycler with following conditions: 95C for 3 minutes, beginning cycling of 98C for 20s, 67C for 20s, and 72C for 3 min. Cycles were allowed to continue on a qPCR machine until reaction neared saturation, as denoted by the exit of exponential phase in amplification. Finally, samples were incubated at 72C for 5min once sufficient cycling occurred. After this reaction, a double-sided SPRI size selection was performed with bounds of 0.5x – 0.7x, where resulting 20 uL eluant was a sequencing-ready library.

### Computational Methods

#### Alignment and generation of cell-gene matrices

The data preprocessing and alignment was performed using a modified SPLiT-seq pipeline (*1*), where the cDNA reads were mapped to either a combined B. S. – E. C genome (ASM49748v1.44 and ASM80076v1.37 from EnsemblBacteria (*2*)) or the *B. subtilis* 168 genome (ASM904v1.45 from EnsemblBacteria) using STAR with the splicing isoform detection switched off (*3*). We chose to use the parent strain *B. subtilis* 168 genome for the growth curve experiment since the respective annotation file had gene names, which the PY79 strain annotation file lacked (using non-standard accession numbers instead). However, we note that using a different genome and annotation increased the total number of detected genes, largely due to the different annotation. In addition, we kept the multimapping reads, assigning a fractional count based on the number of alignments, since bacterial genomes are known to contain overlapping CDSs. We then generated a matrix of gene counts for each cell (N x K matrix, with N cells and K genes).

#### Processing of data from the heat shock experiment

Clustering and data analysis for the species-mixing experiment with heat shock treatment was performed using Scanpy (*4*). We only kept transcriptomes with the number of total reads higher than 200. Then, we removed the ribosomal and tRNA reads from the data, retaining only reads representing the mRNA counts for both species. We further filtered cells based on the mRNA counts, retaining cells expressing >100 reads and >100 genes, and additionally filtered the genes retaining the genes expressed in >5 cells. We then applied standard Scanpy normalization and scaling, dimensionality reduction, and clustering as described in the Scanpy tutorial (*4*, *5*) and below, using 9 PCA components and 45 neighbors for computing the neighborhood graph of cells. For plotting the heat map of the top 6 enriched genes for the heat shock experiment clusters (**Figure S4B**), we additionally filtered the genes by keeping only the genes expressed in at least 40% of the cluster, with a minimum fold change of 2 and in at most 30% of the rest of the data. For making the half-volcano plot of enriched genes in the *E. coli* heat shock cluster (**Figure 1H**), for better clarity we omitted the outlier norR gene with a very high fold change but low significance.

#### Processing of B. S. data from the growth in rich media experiment

Clustering and data analysis for the combined 10 samples of B.S. grown in rich medium was performed using Scanpy (*4*) and verified with Seurat v3 (*6*) and UNCURL (*7*). Experiment 1 sampled OD points 0.5, 1.0, 1.7, 2.0, 2.8, and 3.2, while in experiment 2 we collected OD points 0.5, 1.0, 1.3, 1.7, 2.8, 3.5, 5.3, and 6.0. For the data from both experiments separately, we discarded any transcriptomes with the number of total reads fewer than 200. Then, we filtered the data and retained only reads representing the mRNA counts (excluding the ribosomal and tRNA reads). Finally, we combined the data matrices together. Since the read depth decreased for the higher OD samples (**Figure S15**), for selecting the highest quality data, we implemented differential thresholds for each OD in the combined matrix, retaining top 75% of the cells by read counts for each OD sample. This resulted in retention of 25,214 transcriptomes from both experiments. Finally, we performed batch correction through Scanpy, using a python implementation of ComBat (*8*, *9*).

Cells that passed the QC were clustered using a pipeline described in previous studies (*1*, *5*). We chose to omit the high variance gene selection since the number of genes we detected (on the order of 3,000) was much lower than what is typically detected for mammalian cells and permitted analysis without confining the gene set. In addition, restriction to high variance genes did not result in a major change in the final clustering output. We followed with standard Scanpy normalization and scaling, dimensionality reduction, and clustering (*4*, *5*). Briefly, we computed the neighborhood graph of cells using the top 16 PCA components of the data matrix (n-neighbors = 8). We then used Louvain graph-clustering method to produce the global clustering of the data. For the two-dimensional embedding of the data, we chose to use the t-distributed Stochastic Neighbor Embedding (t-SNE) using a *scikit-learn* implementation (*10*) of the Barnes-Hut t-SNE algorithm (*11*). Top enriched genes for each cluster were computed by Wilcoxon rank-sum test with Benjamini-Hochberg correction.

Sub-clustering of the late OD samples

For finding the finer grain structure within the late OD point data, we took filtered read matrices from cells belonging to OD 5.3 and 6.0 groups and re-run the processing pipeline as described above. Because of the smaller numbers of cells, we chose to use the UMAP algorithm (*12*) to embed the data.

Calculation of Sigma Factors and Transcriptional Regulators Activity

To generate the profiles of activities of sigma factors and transcriptional regulators (TR), we used the *Subti*Wiki resource (*13*). For the TR analysis, each regulon was divided into “positive” and “negative” subregulons based on the mode of regulation (we omitted the rare more complex interactions and focused on the more straightforward transcriptional activation or repression), and we then calculated the average expression of genes within each subregulon for each cluster.

**Supplementary Text**

*B.subtilis* growth curve fit

For the growth curve experiment, we sampled 10 OD points across two experiments. We retained all optical density measurements, however, the time data from the first experiment was not recorded. We fit the growth curve in Figure 2 to the time-stamped OD samples from the second experiment using the formula for a sigmoidal curve: $L/(1+\exp(-k(t-t0)))$. To plot the OD points from the first experiment, we set the fit to the recorded OD curve and solved for time.

TCA cycle gene expression

In cluster 7, in contrast to clusters 3 and 4, instead of excretion we now observe uptake of lactate via *lutP* and conversion to pyruvate by *lutAC* (*14*). The conversion of pyruvate to oxaloacetate is also upregulated by increased expression of *pycA* (*15*). In addition, acetoin, a product of acetate metabolism, in this cluster is actively converted into butanediol by action of *bdhA* (*16*). Cells in this cluster also express more *dctP* implicated in direct import of TCA cycle intermediates succinate, fumarate, malate and oxaloacetate (*17*). In cluster 8, we additionally find enrichment of *pckA* converting oxaloacetate to PEP (**Figures 3C, S8-S9**) (*18*).

PBSX prophage cluster gene expression

Two of the host genes enriched in cluster 13, *mcpC* and *amtB*, are linked to the GlnR regulon (*19*, *20*) which responds to excess nitrogen for nitrogen assimilation. Given that the cells in this cluster were largely in stationary phase and therefore in a nutrient limited environment, it is unlikely that there was excess nitrogen to induce this regulon. It is possible that induction of PBSX might lead to expression of the GlnR host regulon, but further research is needed. A similar hypothesis can be formulated for the expression of *sacA*, which is involved in carbohydrate uptake (*21*, *22*). The other genes identified, *liaL*, *ykuG* (syn. *fadG*), and *recX*, have been shown to be involved in the LiaRS membrane damage response, fatty acid degradation response, and homologous recombination respectively (*23*–*25*).

Competence cluster gene expression

Notably, we detected enrichment of *ywfM* and *hemQ* in the competence cluster. *ywfM* whose product is unknown is located very close but in an opposite orientation to *hemQ* encoding a coproheme decarboxylase implicated in heme biosynthesis. It was prior hypothesized that ComK could be activating divergent transcription by binding to the palindromic site between *yfmK* and *hemQ*, however, *yfmK* and *yfmM* were not previously reported to be co-transcribed. In a similar vein, we detected enrichment of *nin* and *tlpC* with the latter previously not reported to be ComK-induced, however, it is chromosomally located next to the *hxlA/hxlR* gene pair on the opposite strands of DNA divergently transcribed by ComK binding between these genes and activating transcription in both directions. *tlpC* encodes an orphan membrane-bound chemotaxis receptor (*26*). ComK is known to induce expression of chemotaxis proteins, encoded by the genes *flgL,K,M* in the comFA operon which we also detected in our competence cluster. However, competence phenotype is considered suppressive to motility via action of anti-sigma factor *flgM*. Currently the exact ligand for *tlpC* is not known except *tlpC* being functionally connected to *A. thaliana* root colonization (*27*) and it is therefore hypothesized to be involved in other cellular process involving a two-component signal transduction system (*26*). *hemQ* product, coproheme decarboxylase, on the other hand, functions in heme biosynthesis. Various intermediary metabolism genes are known to be upregulated by ComK but they were not previously known to include the heme biosynthetic pathway. Finally, another gene not hitherto connected to K-state but enriched in the competence cluster is *trmF*, a folate- and FAD-dependent tRNA methyltransferase involved in tRNA maturation. This finding is in line with another methyltransferase *gidB* acting on 16S rRNA which has been found enriched in competent cells previously (*28*).

**Supplementary Figures**

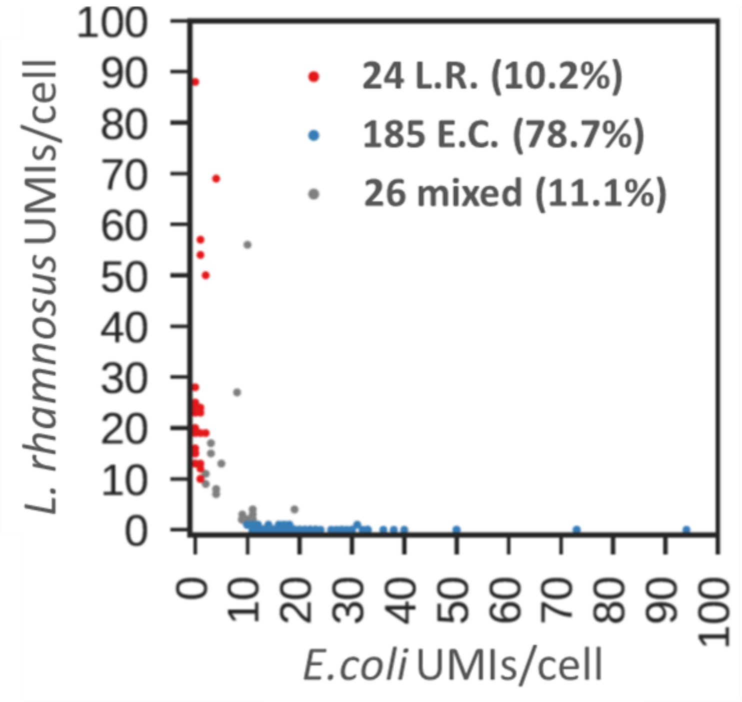


**Figure S1. Direct application of SPLiT-seq to mixed *Lactobacillus rhamnosus* and *E. coli* cells.**

A barnyard plot showing each putative single-cell transcriptome as a dot aligned to the combined *L. rhamnosus* –*E. coli* (E.C.) genome. The cells with <10 reads were filtered out. UMI – unique molecular identifier.

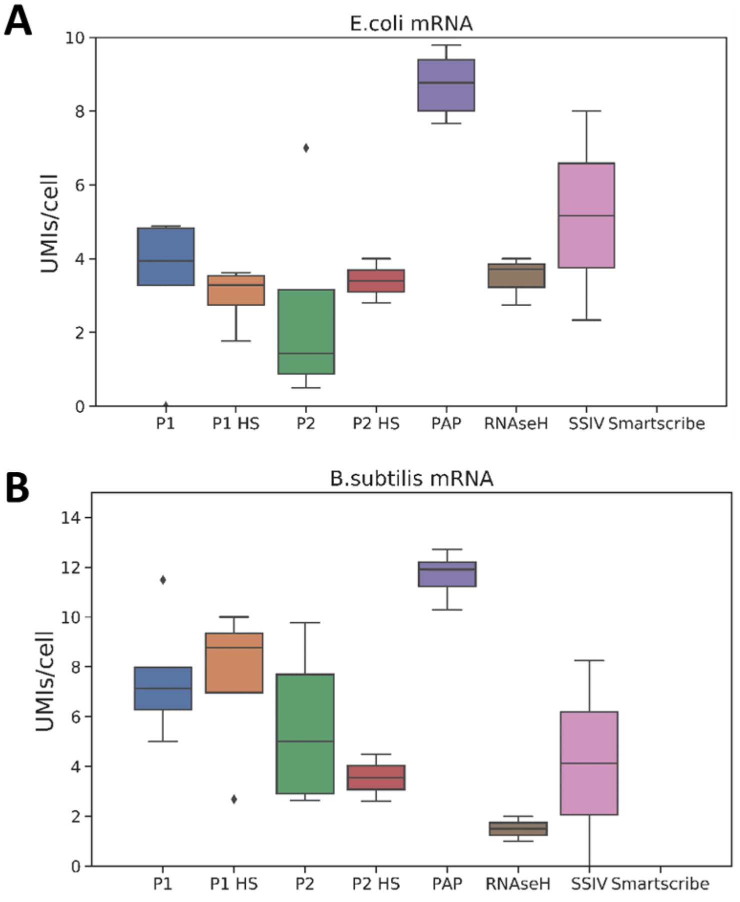


**Figure S2. Optimization of μSPliT permeabilization and mRNA enrichment steps.** Sample test of differential conditions for permeabilization and mRNA enrichment for **(A)** *E. coli* and **(B)** *B. subtilis* read counts. UMI – unique molecular identifier. Conditions tested: "P1" - 0.5%Triton-X100 + Lysozyme (5mg\ml) for both species; "P2" - 0.5%Triton-X100 for *E. coli* and 0.5%Triton-X100 + Lysozyme (20mg\ml) for *B. subtilis*; "HS" – heat shocked samples; "PAP" – *E. coli* Poly(A) Polymerase treatment; "RNAseH" – reverse transcription (RT) with rRNA-specific probes followed by RNAseH digestion; "SSIV" – SuperScript IV reverse transcriptase (ThermoFisher) instead of Maxima reverse transcriptase (ThermoFisher); "Smartscribe" – SMARTScribe reverse transcriptase (Takara Bio) instead of Maxima reverse transcriptase (ThermoFisher). For the latter three conditions "P1" permeabilization was used.

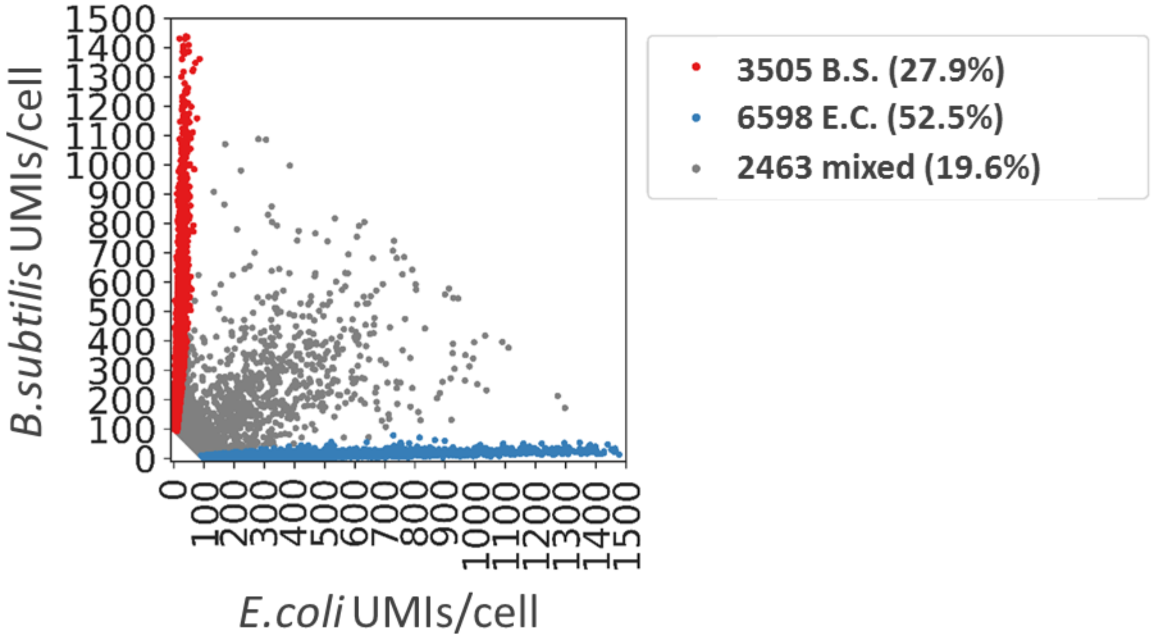


**Figure S3. µSPLiT procedure results in bacterial aggregate formation in absence of sonication or vortexing.**
A barnyard plot showing each putative single-cell transcriptome as a dot aligned to the combined B.S. –E.C. genome. UMI – unique molecular identifier. The cells with <100 reads were filtered out. The mixed cells are expected to originate from the physical aggregates traveling through barcoding reactions together.

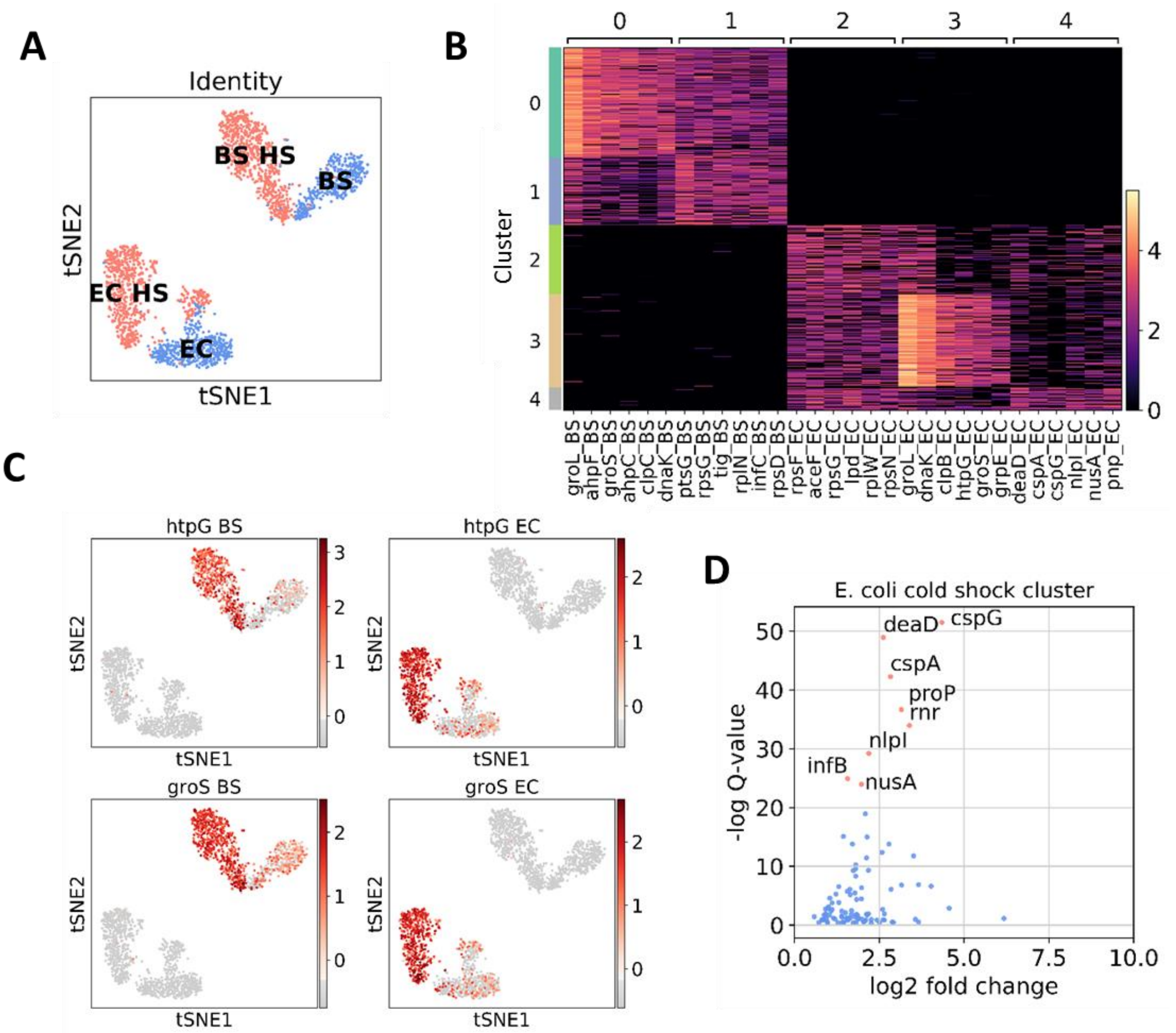


**Figure S4. Transcriptional responses of *E. coli* and *B. subtilis* cells to heat shock. (A)** t-SNE plot showing cluster assignments based on the first barcode label ("HS" indicates heat shocked samples) and the genes expressed in the cluster ("EC" – *E. coli*; "BS" – *B. subtilis*). **(B)** Top 6 genes enriched for each cluster in Figure 1F). **(C)** Expression of select classical heat shock markers for both *E. coli* and *B. subtilis* overlayed on the t-SNE. **(D)** Genes enriched in cluster 4, *E. coli* cold shock cluster.

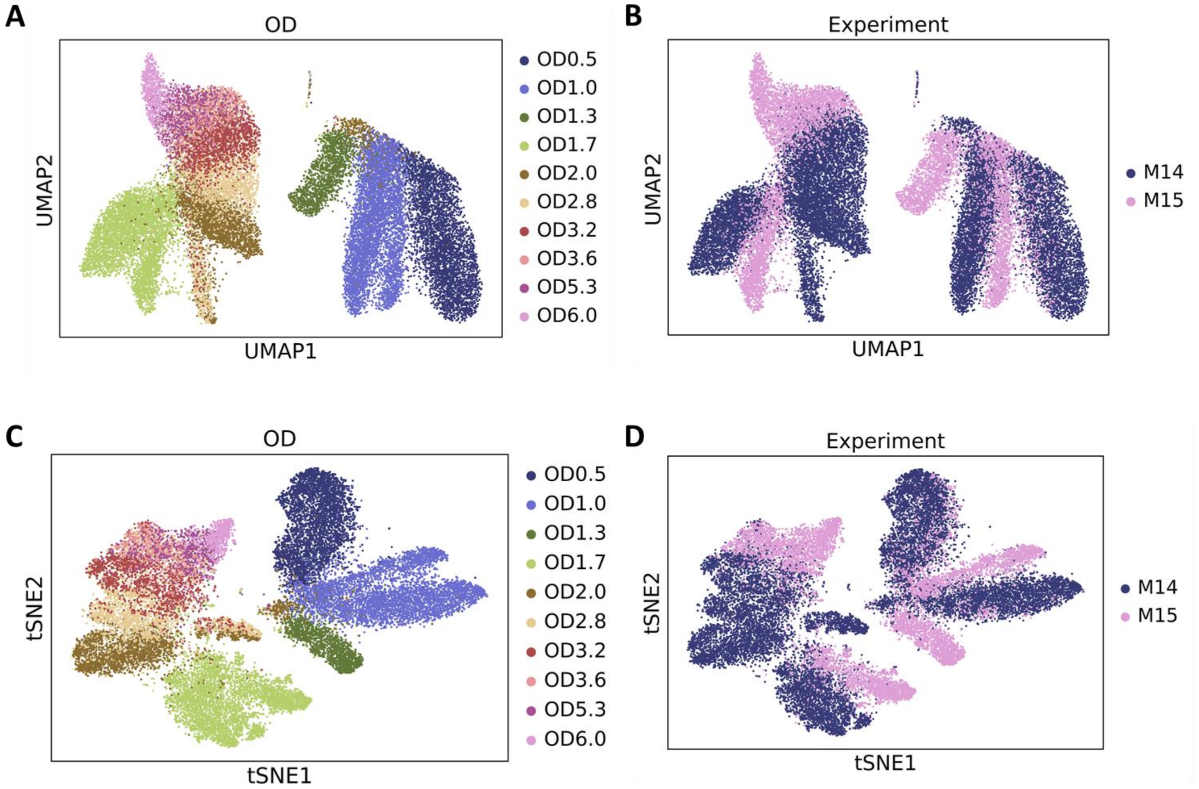


**Figure S5. 25,214 *B. subtilis* cells sampled across the growth curve. (A-B)** UMAP embedding of data from the rich medium growth experiment, colored by either OD label (A) or by experiment (B, "M14" – experiment 1, "M15" – experiment 2). **(C-D)** As in (A-B), but with the t-SNE embedding.

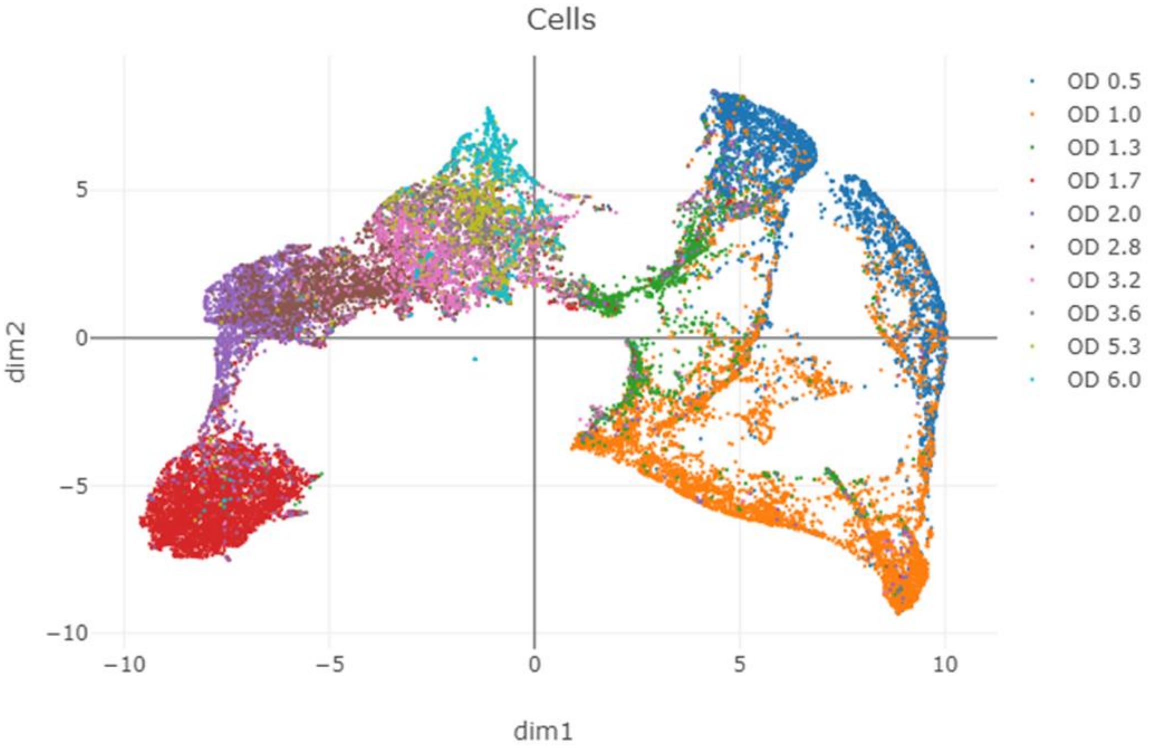


**Figure S6.** UNCURL processing and embedding of data from the rich medium growth experiment, colored by the OD label.

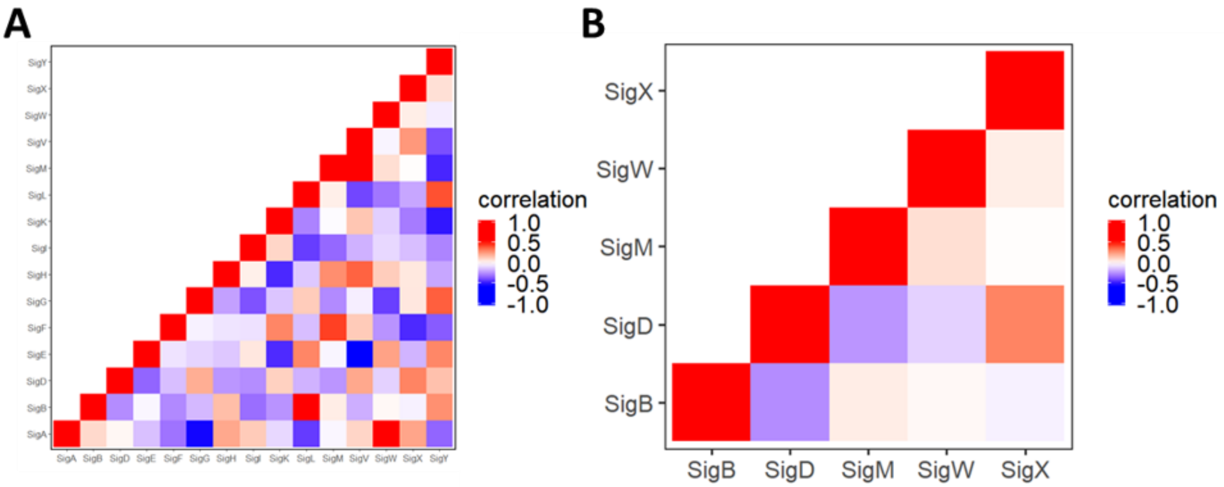


**Figure S7. Expression correlation of genes regulated by alternative sigma factors.** The mean expression for each sigma regulon was calculated and normalized for each of the 25,214 cells and correlated to the mean expression of the other regulons. (**A**) shows a correlation heatmap for the 16 central alternative sigma factors. (**B**) shows the correlations of a select set of sigma factors described in (*29*).

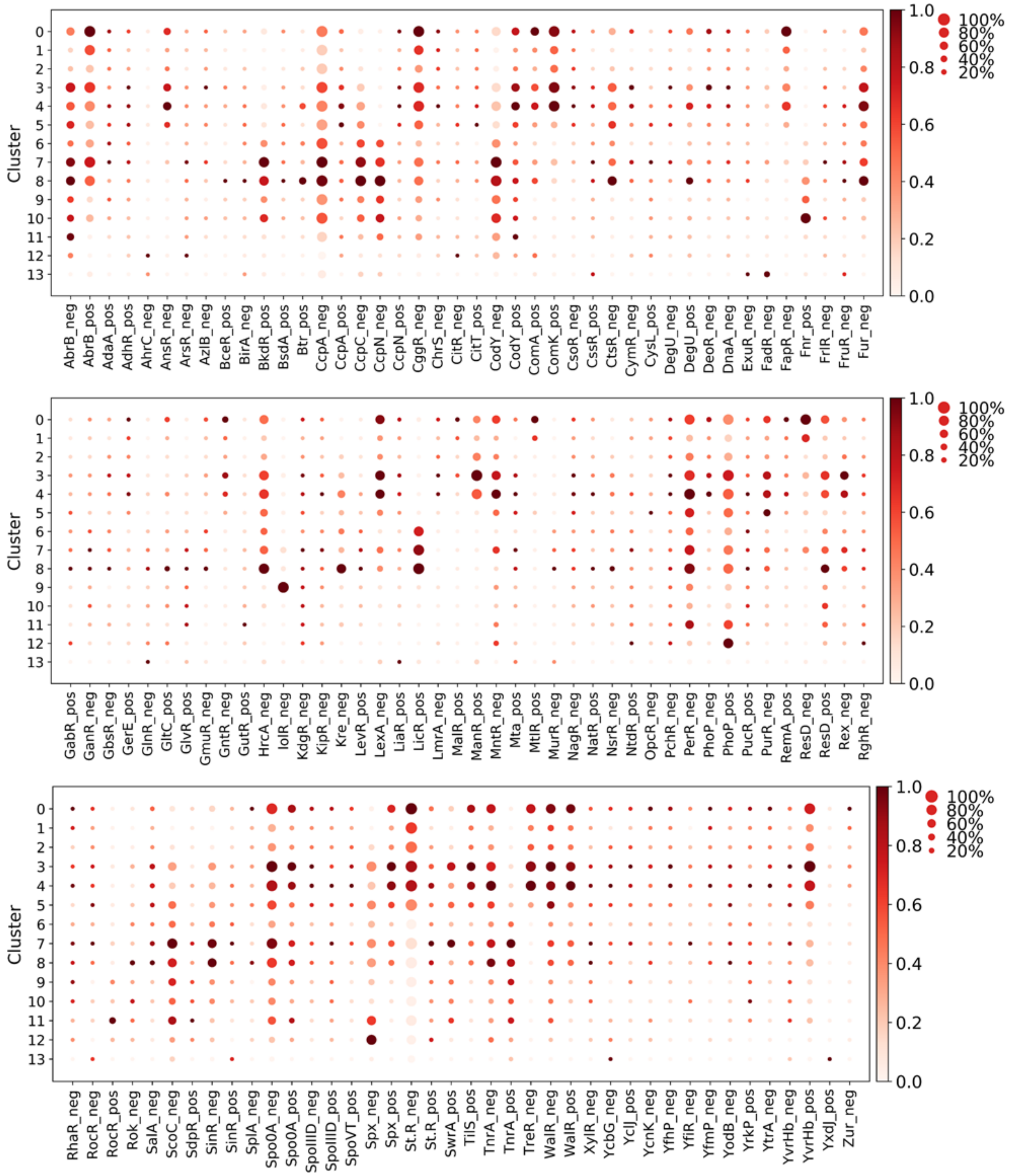


**Figure S8. N**ormalized detected transcriptional regulators (TR) activity per cluster (see **Materials and Methods**). The size of each dot indicates the proportion of cells in the cluster in which the sigma factor is active, while the color indicates the average activity. “Neg” for a given TR indicates that the activity was calculated for the genes in the respective regulon negatively regulated by this TR, and “pos” indicates the activity was calculated for the genes whose expression is positively regulated by the given TR.

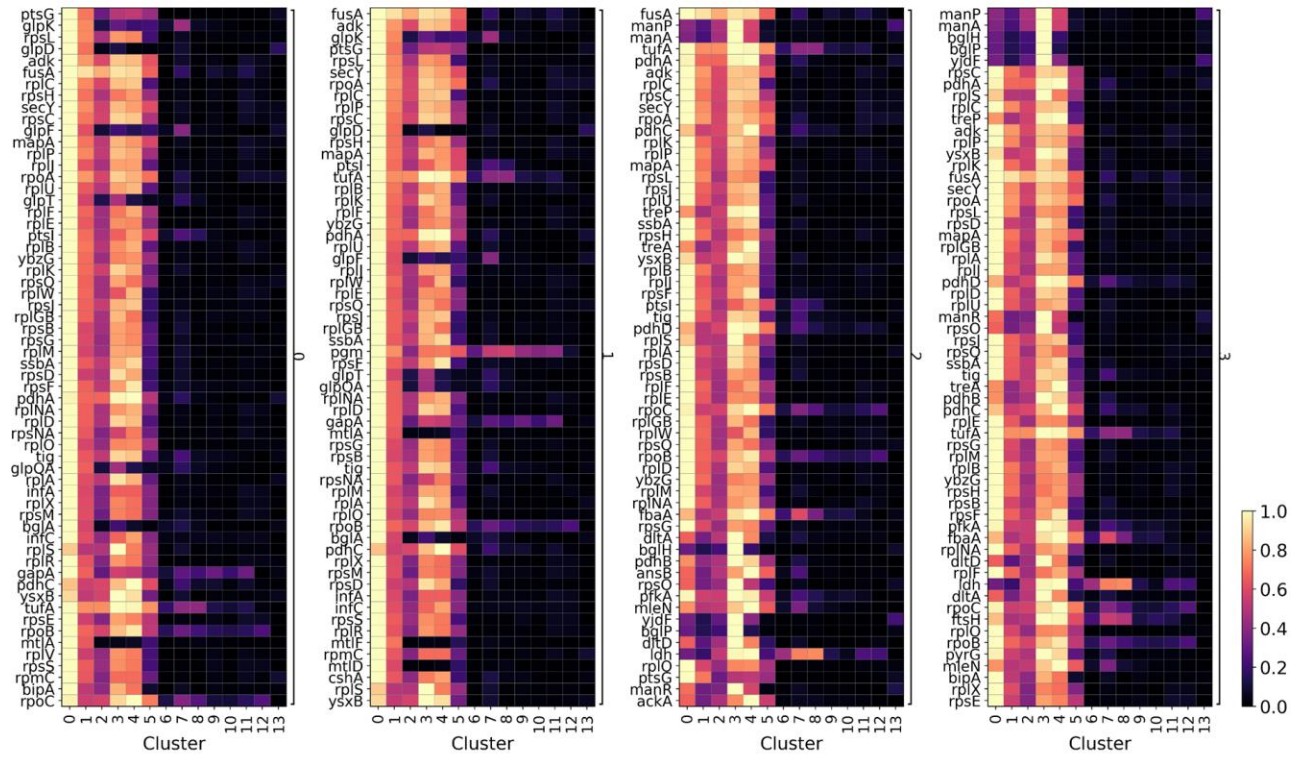


**Figure S9. Top 60 expressed genes for clusters 0-3.**

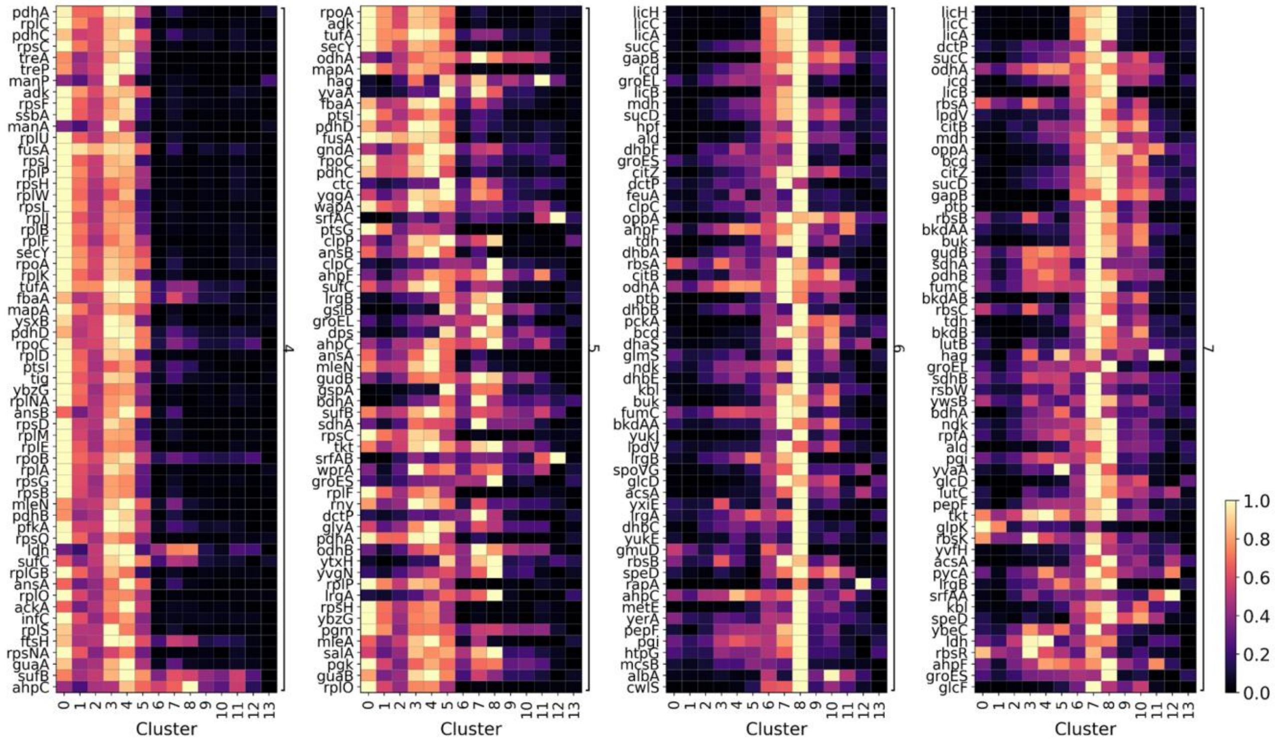


**Figure S10. Top 60 expressed genes for clusters 4-7.**

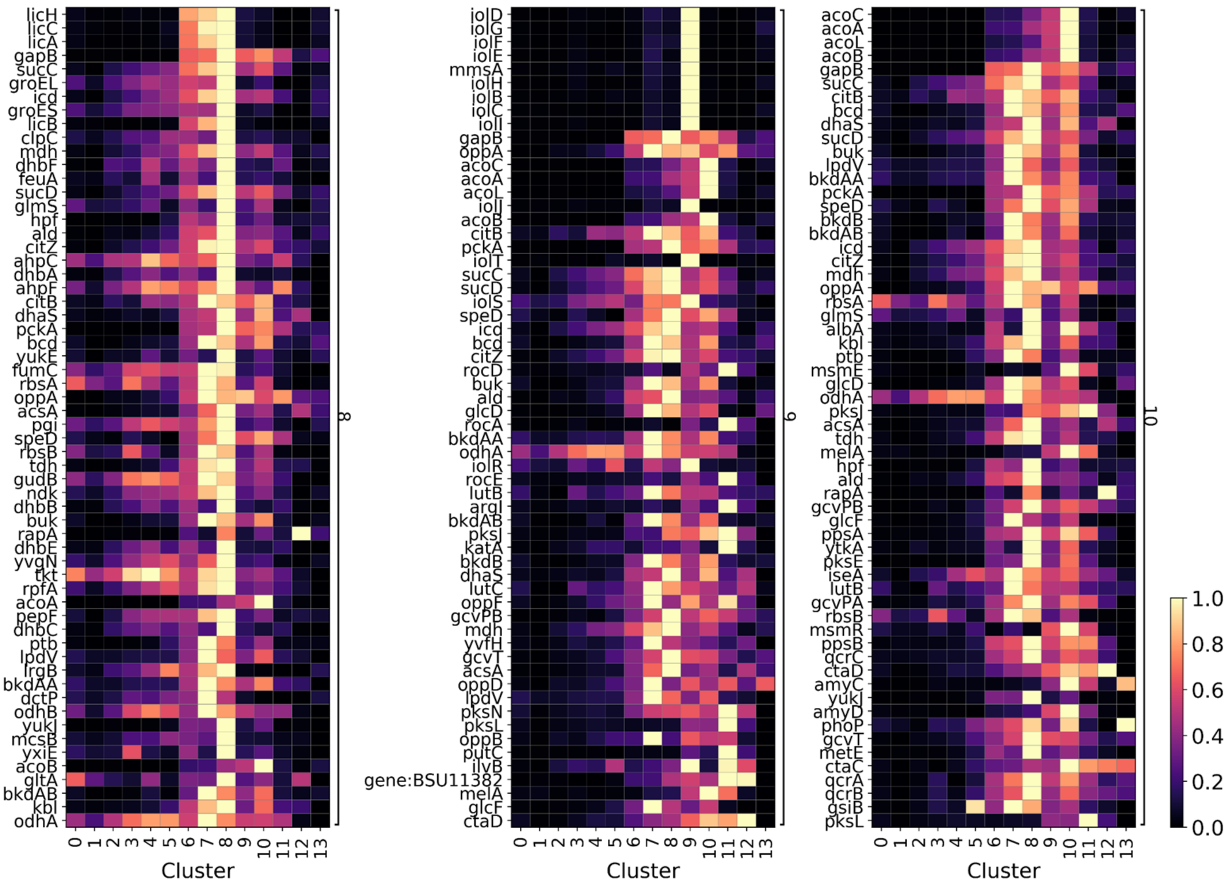


**Figure S11. Top 60 expressed genes for clusters 8-10.**

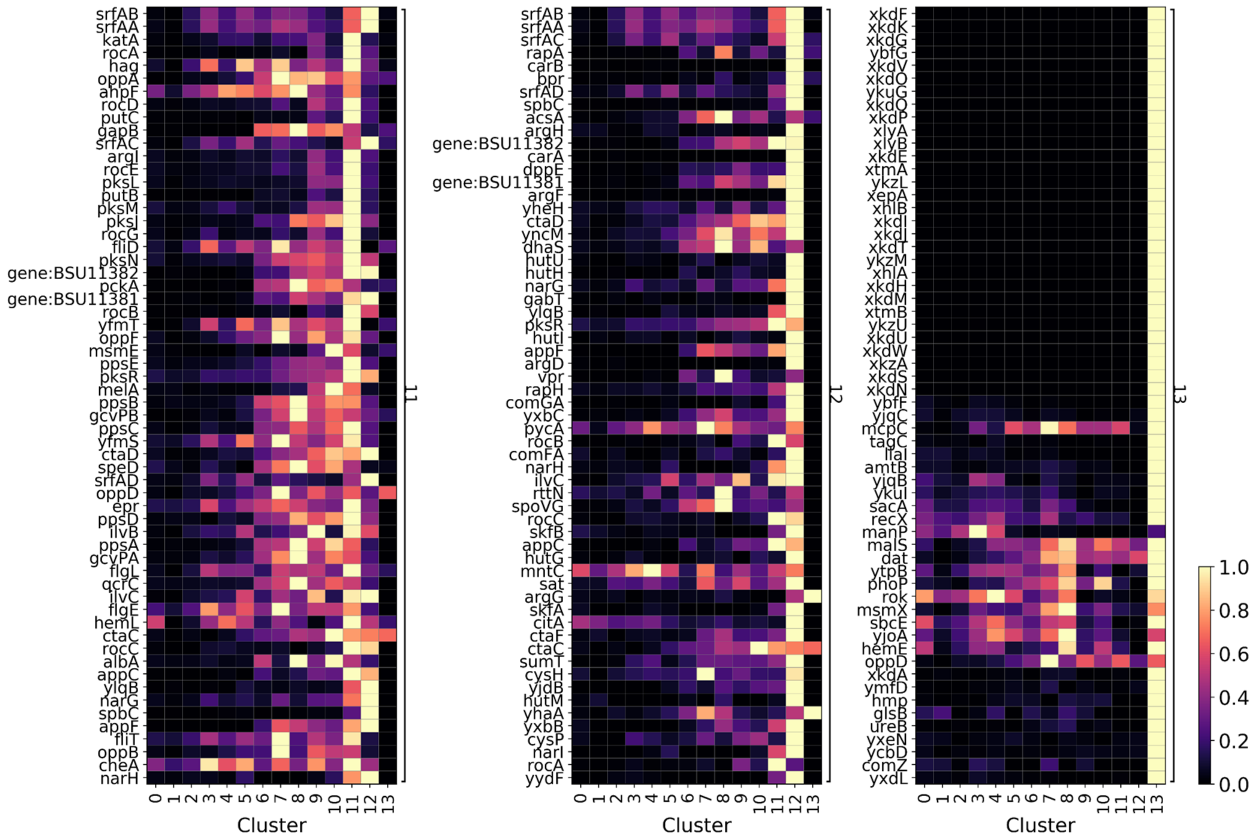

**Figure S12. Top 60 expressed genes for clusters 11-13.**

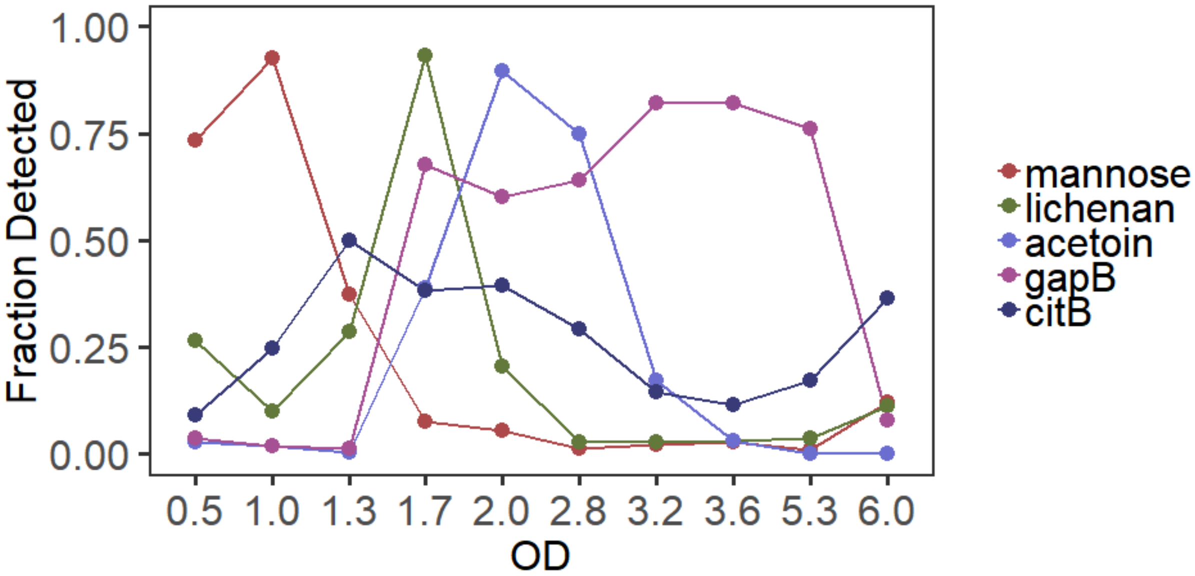


**Figure S13.** Fraction of cells expressing the indicated carbon metabolism genes or operons as a function of OD. The genes used to generate the curves are listed in Supplementary Table S2.



A

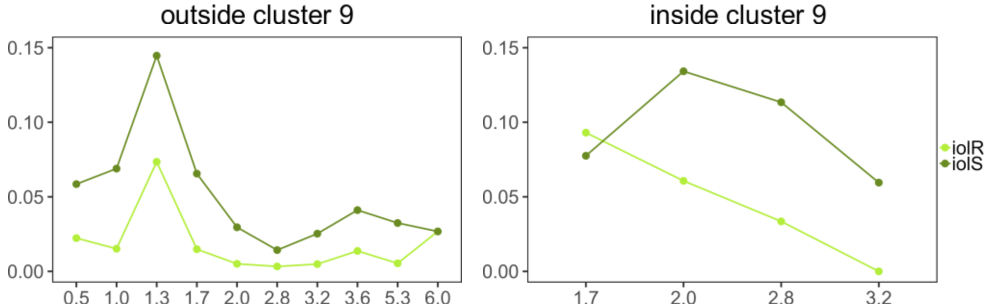

**Figure S14. Fraction of cells detected to express each of the three inositol utilization operons.** (A) iolRS, (B) iolAJ, (C) iolT operon expressing cells fraction plotted for cells either outside or inside cluster 9 across OD measurements. Cells were classified as expressing if we detected at least one transcript from any of the genes in operon. Note that within cluster 9, only ODs 1.7-3.2 are shown as only those samplings had meaningful (>5) numbers of cells included in this cluster.

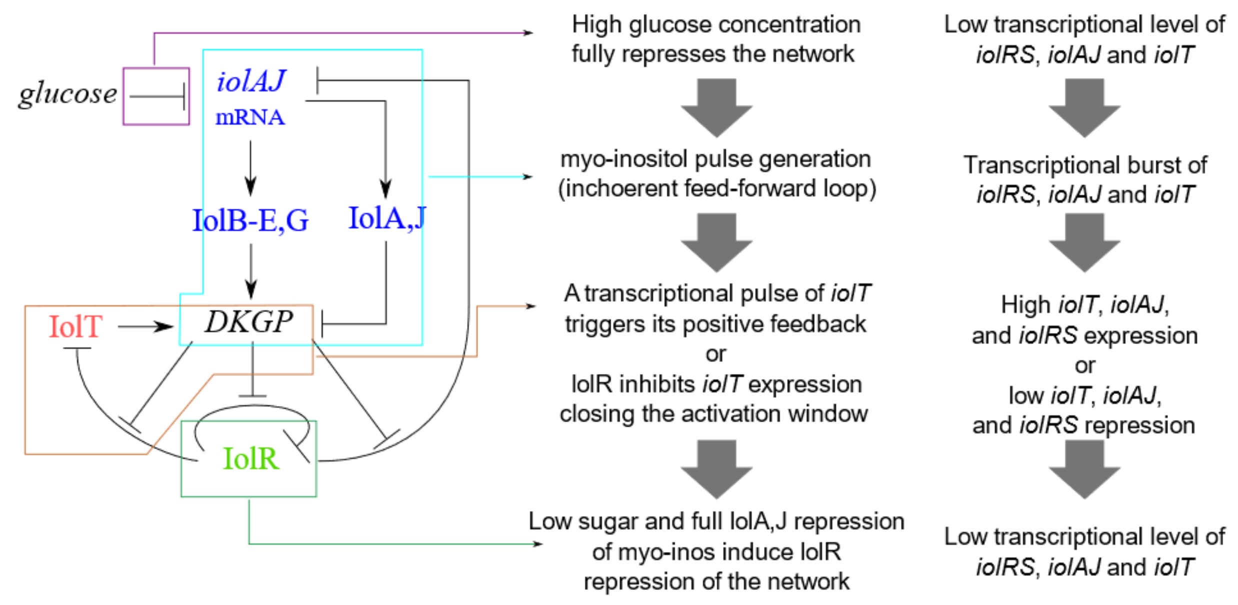


**Figure S15.** Myo-inositol utilization pathway diagram and model description with potential intracellular states that explain the expression heterogeneity we observe.

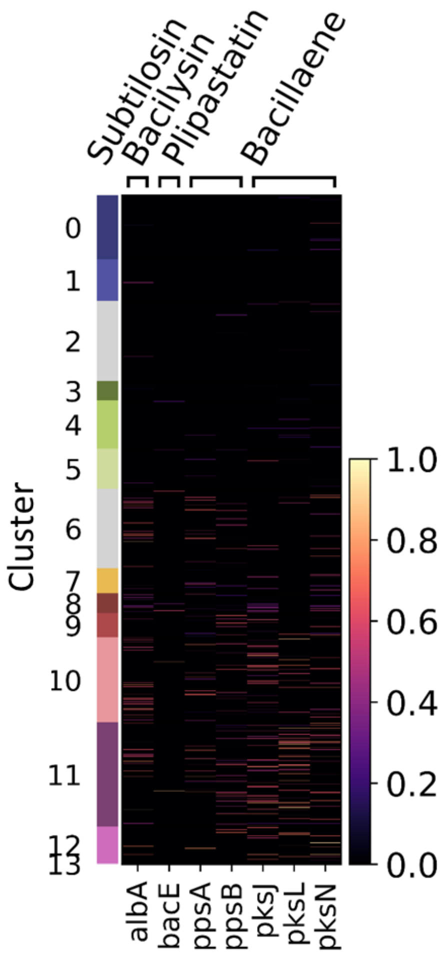


**Figure S16.** Normalized gene expression of the antimicrobial compound production pathways in single cells for each cluster.

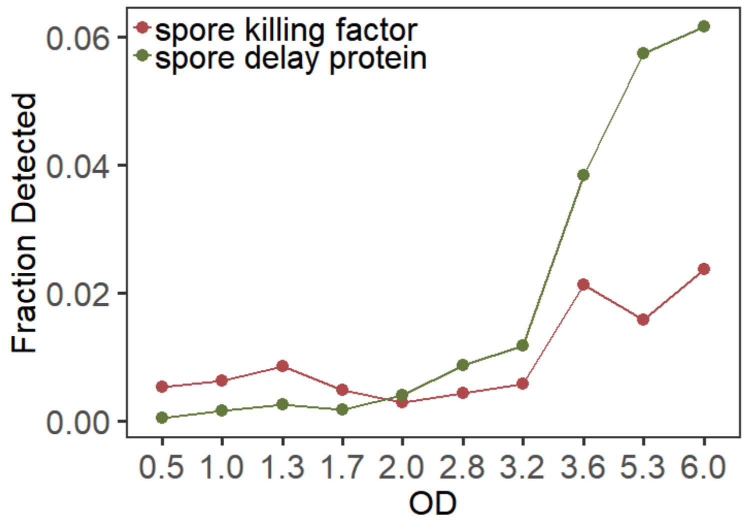


**Figure S17.** Fraction of cells expressing the indicated Skf and Sdp family toxins as a function of OD. The genes used to generate the curves are listed in Supplementary Table S2.

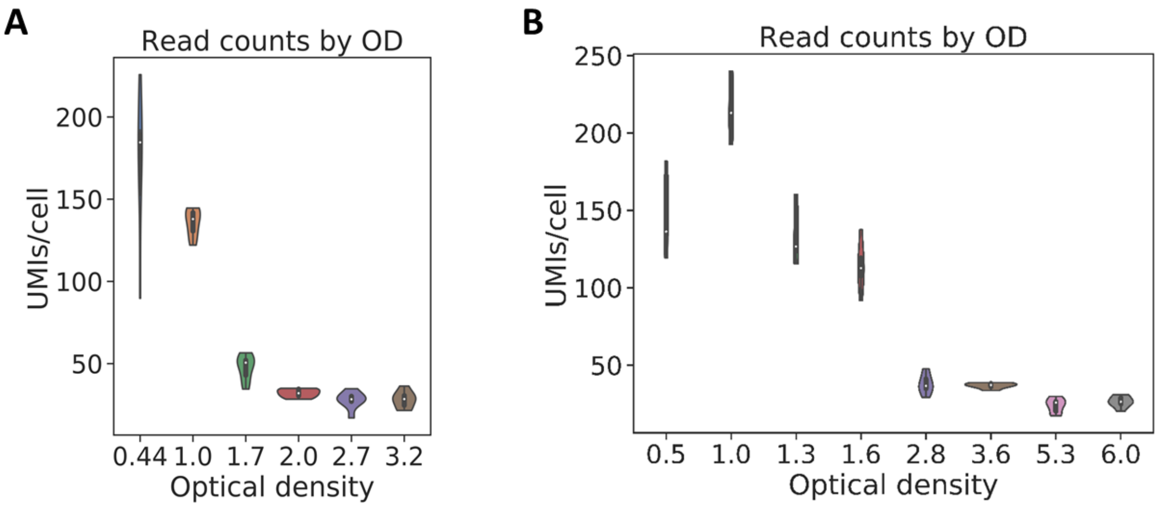


**Figure S18. Read counts for the *B. subtilis* L-broth growth curve experiments. (A)** Experiment 1. **(B)** Experiment 2. UMI – unique molecular identifier.

**Supplementary Tables**

| Fixation | Permeabilization | rRNA depletion | Reverse transcription |
|---|---|---|---|
| 100% Ethanol | 0.5%Triton | Ribosomal RT + RNAseH | Maxima RT + Ficoll |
| 50% Ethanol | Bead beater | TEX Buffer A | Superscript IV RT |
| 4% Formalin + 100% Ethanol | Melittin | TEX Buffer B | Smartscribe RT |
| 4% Formalin + 50% Ethanol | Lysozyme (1mg\ml, 30min) | TEX Buffer A 2x | Maxima RT 50C 15min 60C 15min |
| 1.33% Formalin + 100% Ethanol | Lysozyme (5mg\ml, 30min) | Poly A Polymerase (PAP) + TEX Buffer A | Maxima RT 50C for 30 min |
| 1.33% Formalin | Lysozyme (20mg\ml, 15min) | Poly A Polymerase (PAP) | Maxima RT 50C for 1hr |
| 4% Formalin for 10 min | 0.5%Triton + Lysozyme (20mg\ml, 15min) | | Maxima RT 50C for 4hrs |
| 4% Formalin for 1 hr | 0.5%Tween + Lysozyme (5mg\ml, 15min) | | Maxima RT Cycling RT (SPLiT-seq) |
| 4% Formalin overnight | 0.04%Tween + Lysozyme (2.5mg\ml, 15min) | | Maxima RT 23C 10 min 50C 50 min |

**Table S1. Tested fixation, permeabilization, mRNA enrichment, and RT conditions.**

| FUNCTION | OPERON | GENE | … | FUNCTION | OPERON | GENE |
|---|---|---|---|---|---|---|
| antimicrobial | subtilosin | sboA | | motility | fla-che operon | fliP |
| antimicrobial | subtilosin | sboX | | motility | fla-che operon | fliQ |
| antimicrobial | subtilosin | albA | | motility | fla-che operon | fliR |
| antimicrobial | subtilosin | albB | | motility | fla-che operon | flhB |
| antimicrobial | subtilosin | albC | | motility | fla-che operon | flhA |
| antimicrobial | subtilosin | albD | | motility | fla-che operon | flhF |
| antimicrobial | subtilosin | albE | | motility | fla-che operon | flhG |
| antimicrobial | subtilosin | albF | | motility | fla-che operon | cheB |
| antimicrobial | subtilosin | albG | | motility | fla-che operon | cheA |
| antimicrobial | bacillaene | pksC | | motility | fla-che operon | cheW |
| antimicrobial | bacillaene | pksD | | motility | fla-che operon | cheC |
| antimicrobial | bacillaene | pksE | | motility | fla-che operon | cheD |
| antimicrobial | bacillaene | pksF | | motility | fla-che operon | sigD |
| antimicrobial | bacillaene | pksG | | motility | fla-che operon | swrB |
| antimicrobial | bacillaene | pksH | | stress response | mcs kinase | mcsA |
| antimicrobial | bacillaene | pksI | | stress response | mcs kinase | mcsB |
| antimicrobial | bacillaene | pksJ | | stress response | clpP proteases | clpC |
| antimicrobial | bacillaene | pksL | | stress response | clpP proteases | clpP |
| antimicrobial | bacillaene | pksM | | stress response | clpP proteases | clpE |
| antimicrobial | bacillaene | pksN | | stress response | clpP proteases | clpX |
| antimicrobial | bacillaene | pksR | | stress response | chaperonin | groEL |
| antimicrobial | bacillaene | pksS | | stress response | chaperonin | groES |
| antimicrobial | bacilysin | bacB | | metal import | manganese uptake | mntA |
| antimicrobial | bacilysin | bacC | | metal import | manganese uptake | mntB |
| antimicrobial | bacilysin | bacD | | metal import | manganese uptake | mntC |
| antimicrobial | bacilysin | bacE | | metal import | manganese uptake | mntD |
| antimicrobial | bacilysin | bacH | | metal import | manganese uptake | mntH |
| antimicrobial | plipastatin | ppsA | | metal import | bacillibactin | dhbA |
| antimicrobial | plipastatin | ppsB | | metal import | bacillibactin | dhbC |
| antimicrobial | plipastatin | ppsC | | metal import | bacillibactin | dhbE |
| antimicrobial | plipastatin | ppsD | | metal import | bacillibactin | dhbB |
| antimicrobial | plipastatin | ppsE | | metal import | bacillibactin | dhbF |
| antimicrobial | endoA/I | ndoA | | metal import | siderophore uptake | feuA |
| antimicrobial | endoA/I | ndoAI | | metal import | siderophore uptake | feuB |
| antimicrobial | spore killing factor | skfA | | metal import | siderophore uptake | feuC |
| antimicrobial | spore killing | skfB | | metal import | siderophore | fhuB |

| | | | | | | |
|---|---|---|---|---|---|---|
| | factor | | | | uptake | |
| antimicrobial | spore killing factor | skfE | | metal import | siderophore uptake | fhuC |
| antimicrobial | spore killing factor | skfF | | metal import | siderophore uptake | fhuD |
| antimicrobial | spore killing factor | skfG | | metal import | siderophore uptake | fhuG |
| antimicrobial | spore delay protein | sdpB | | metal import | siderophore uptake | yxeB |
| antimicrobial | spore delay protein | spbC | | carbon metabolism | lichenan | licA |
| motility | surfactin | srfAA | | carbon metabolism | lichenan | licB |
| motility | surfactin | srfAB | | carbon metabolism | lichenan | licC |
| motility | surfactin | srfAC | | carbon metabolism | lichenan | licH |
| motility | surfactin | srfAD | | carbon metabolism | acetoin | acoA |
| motility | flagellin protein (hag) | hag | | carbon metabolism | acetoin | acoB |
| motility | fla-che operon | flgB | | carbon metabolism | acetoin | acoC |
| motility | fla-che operon | flgC | | carbon metabolism | acetoin | acoL |
| motility | fla-che operon | fliE | | carbon metabolism | mannose | manA |
| motility | fla-che operon | fliF | | carbon metabolism | mannose | manP |
| motility | fla-che operon | fliG | | carbon metabolism | gapB | gapB |
| motility | fla-che operon | fliH | | carbon metabolism | citB | citB |
| motility | fla-che operon | fliI | | carbon metabolism | iolAJ | iolA |
| motility | fla-che operon | fliJ | | carbon metabolism | iolAJ | iolB |
| motility | fla-che operon | ylxF | | carbon metabolism | iolAJ | iolC |
| motility | fla-che operon | fliK | | carbon metabolism | iolAJ | iolD |
| motility | fla-che operon | flgD | | carbon metabolism | iolAJ | iolE |
| motility | fla-che operon | flgE | | carbon metabolism | iolAJ | iolF |
| motility | fla-che operon | fliL | | carbon metabolism | iolAJ | iolG |
| motility | fla-che operon | fliM | | carbon metabolism | iolAJ | iolH |

| | | | | | | |
|---|---|---|---|---|---|---|
| motility | fla-che operon | fliY | | carbon metabolism | iolAJ | iolI |
| motility | fla-che operon | cheY | | carbon metabolism | iolAJ | iolJ |
| motility | fla-che operon | fliZ | | carbon metabolism | iolRS | iolR |
| | | | | carbon metabolism | iolRS | iolS |
| | | | | carbon metabolism | iolT | iolT |

**Table S2. Genes and operons used in select carbon metabolism and lifestyle analyses.**
